# A novel phenylpropanoid methyl esterase enables catabolism of aromatic compounds that inhibit biological nitrification

**DOI:** 10.1101/2023.06.02.543320

**Authors:** Andrew Wilson, Elise Van Fossen, Ritu Shrestha, Valentine Trotter, Brenton Poirier, Andrew Frank, William Nelson, Young-Mo Kim, Adam Deutschbauer, Robert Egbert, Joshua R. Elmore

## Abstract

Agriculture is the largest source of greenhouse gases (GHG) production. Conversion of nitrogen fertilizers into more reduced forms by microbes through a process known as biological nitrification drives GHG production, enhances proliferation of toxic algal blooms, and increases cost of crop production. Some plants reduce biological nitrification in soils by exuding a diverse array of biological nitrification inhibitors (BNIs) that inhibit the ammonium oxidizing microbes responsible for nitrification. Applying synthetic biology to enhance and transfer BNI production into food and bioenergy crops is a promising approach to reduce nitrification but the success of this strategy is contingent upon improving our limited understanding of BNIs mechanisms-of-action and degradation in the soil. We addressed this gap by investigating the previously unknown metabolic route through which a prominent class of aromatic BNIs known as phenylpropanoid methyl esters (PPMEs) are catabolized. While neither transcriptomics (RNAseq) or high-throughput functional genomics (RB-TnSeq) methods reduced the genetic search space for pathway discovery into a tenable number of genes, the combination of narrowed the search space to a collection of 4 proteins of unknown function. Using genetic and biochemical analyses we found that two previously uncharacterized enzymes, including a novel phenylpropanoid methyl esterase, funnel PPMEs into an established phenylpropanoid catabolism pathway. Transfer of these two enzymes into bacteria capable of using other phenylpropanoids use of PPMEs as carbon sources. This work both provided insight into BNI catabolism and is the first step towards development of model *in vivo* plant-microbe systems for studying BNI mechanisms under well controlled conditions.

## Introduction

Agriculture has become the largest source of greenhouse gases (GHG) on the planet^1^. While application of N-fertilizers increases crop yields^2^, ∼70% of applied N-fertilizer is ultimately lost to the atmosphere as NO_2_, a potent GHG, or to waterways as NO^3-^, which enables toxic algal blooms (**Fig. 1**)^3,4^. Nitrification is the process by which these deleterious compounds are formed ^5^. Thus, reducing nitrification is not only critical to achieving the emissions reductions required to limit climate change, but also for abolishing algal blooms and NO_x_ emissions that reduce air quality.

**Fig. 1.**
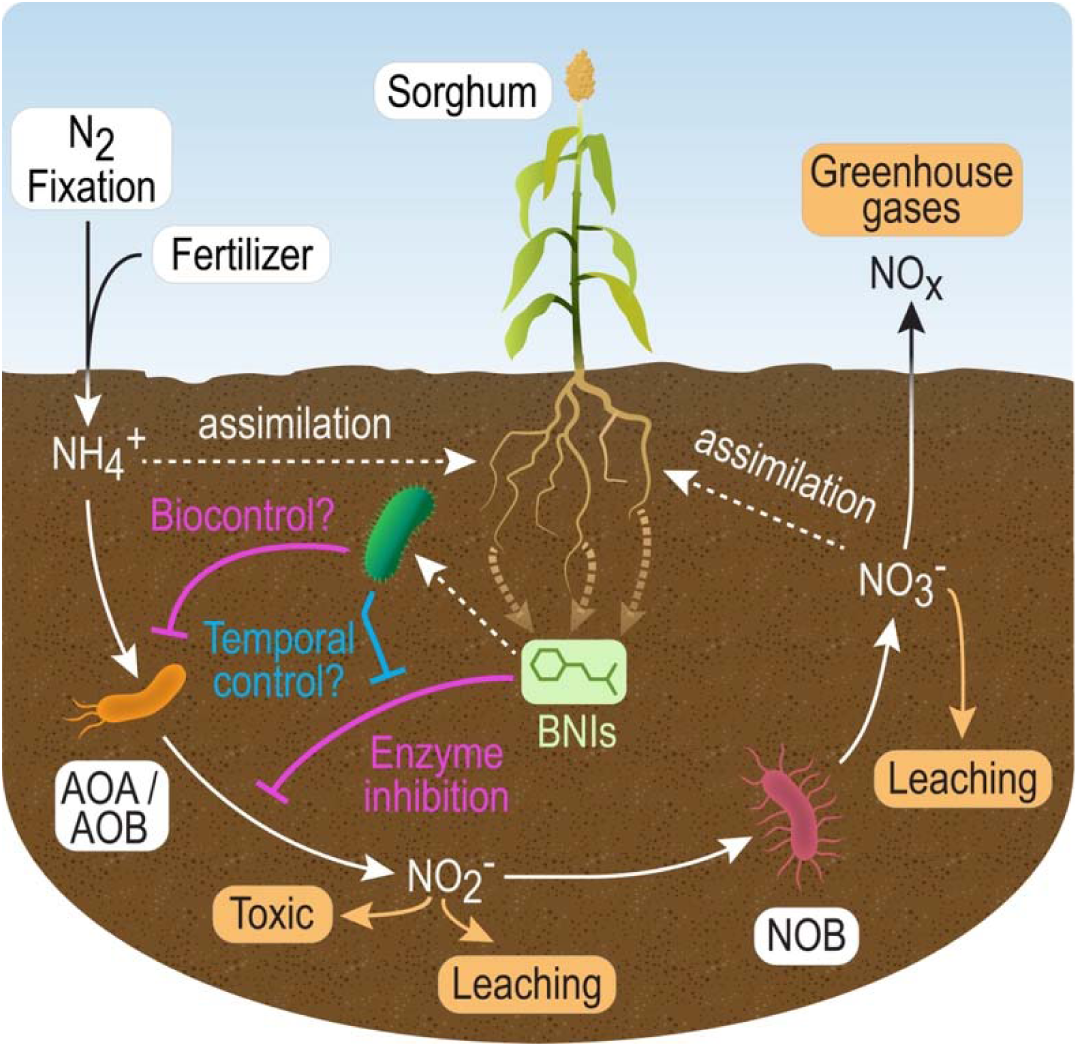
Biological inhibition of the nitrification cycle. Ammonium oxidizing archaea (AOA) and bacteria (AOB) convert excess N in the soil into forms that lead out of soils and are associated with toxicity. Plants exude biological nitrification inhibitor chemicals (BNIs) through their roots to prevent loss of soil nitrogen through this process. Our understanding of BNI modes-of-action are limited by existing experimental methods which focus on enzyme inhibition. A more complete understanding of the nitrogen cycle will require we investigate the microbes enriched by and capable of consuming BNIs and assessing their role in the process. For example, enriched microbes may enhance inhibition through biocontrol of AOA/AOB growth or exert temporal control over enzyme inhibition through mineralization of BNI compounds.

Nitrification, the conversion of reduced forms of nitrogen (NH_3,_ NH_4_^+^) into nitrate (NO_3_^-^), is largely a consequence of microbial oxidation, and reducing this microbial activity in soils is essential to limit N-pollution^6^. Multiple approaches are either currently used or under active development to reduce biological nitrification. One approach used by the agriculture industry is the co-application of chemical nitrification inhibitors with fertilizers^7,8^. Inhibitor chemicals are selected based upon their ability to inhibit activity of the highly conserved ammonium oxidase (AMO) and hydroxylamine oxidoreductase (HAO) enzymes that perform biological nitrification. These enzymatic functions are limited to a small collection of archaeal (AOA) and bacterial (AOB) lineages, and blocking these activities abolishes their ability to grow. Another approach is to limit the application of chemical fertilizers. For this, researchers and several companies (e.g., Pivot Bio) are developing engineered microbial and plant^9^ systems that can be deployed in agricultural soils to enhance fixation of elemental nitrogen (N_2_) into biologically available nitrogen (NH_3_)^9–12^. Supplementing soils with these engineered N_2_-fixing bacteria can substantially reduce the amount of chemical N-fertilizers required for robust crop yields. While effective, these two approaches alone will be insufficient to address the complex challenge of reducing nitrification.

Enhancing biological inhibition of nitrification by plant-produced biological nitrification inhibitors (BNIs) is another highly promising method of reducing nitrification and enhancing plant nitrogen use efficiency^13^. An increasing number of plants, including the food and potential bioenergy crop *Sorghum bicolor*, are known to exude chemicals that inhibit biological nitrification through their roots^14–16^. A diverse array of compounds, including phenolics, lipophilic benzoquinones, terpenes, and diols function as BNIs^15,16^. Advances in synthetic biology have enabled microbe and plant engineering for production of chemicals similar to many BNIs (e.g., *p-*coumaric acid, terpenes)^17–19^. Thus, a promising third approach is to engineer organisms that synthesize and secrete BNIs into the rhizospheres of plants that do not produce them naturally.

However, for such a synthetic biology approach to reducing nitrification to succeed, we must have more information regarding BNI modes-of-action, knowledge of BNI metabolism, and simplified experimental methods to evaluate engineered systems *in situ*. Currently, BNIs are primarily identified using *in vitro* enzyme activity assays^20^ and many are thought to exclusively function by their ability inhibit the activity of AMO or HAO enzymes. The diversity of BNI compounds and knowledge of these compounds other functions in the environment^21,22^ (e.g., herbicidal activity of the BNI sorgoleone^22^) suggest the reality is more complicated^16^. Plants are known to respond to various stress conditions by exuding compounds into the rhizosphere that encourage growth of microbes with specific beneficial functions (e.g., N-fixation, anti-fungal activity)^23–25^. Similarly, exuded BNIs may also perform secondary BNI functions, such as encouraging growth of specific microbes that directly or indirectly inhibit growth of ammonium oxidizing microbes (**Fig. 1**).

Several tasks must be performed before BNIs can be comprehensively evaluated for alternate functions. First, we must identify and isolate microbes that are enriched by BNI compounds. Second, we must characterize the metabolic pathways associated with either BNI catabolism or modification. Third, we must generate mutant microbes lacking these pathways. Comparing BNI capabilities of these enriched microbes with and without the associated pathways will be critical for controlled experiments that directly test the impact of the bacterium, rather than the compound, upon biological nitrification. As a first step towards this, we perform the above steps for several phenylpropanoid methyl esters (PPMEs), a class of phenolic BNIs produced by multiple plants^14,21,26^. Prior to this work, the pathways for PPME catabolism were unknown. Here we characterize the pathway used to catabolize PPMEs and identify a novel enzyme family that performs the first step in PPME catabolism. Finally, we show this metabolic pathway can be transferred into multiple heterologous hosts.

## Results

### Screening bacteria for the ability to utilize phenylpropanoid methyl esters as a sole carbon source

Phenylpropanoid methyl esters (PPME) function as biological nitrification inhibitors^14,26^, but the mechanisms of through which they inhibit biological nitrification and how they are ultimately removed from the environment are not fully understood. Pseudomonads are ubiquitous in soil and rhizosphere environments, where they frequently encounter phenylpropanoids (e.g., *p-*coumaric acid and ferulic acid) from plant root exudates and decomposing plant biomass. Many environmental Pseudomonads have been isolated with or shown to utilize phenylpropanoid carbon sources^27–29^, and thus may also be able to utilize PPMEs as carbon sources.

To identify organisms to capable of using these compounds as carbon sources we evaluated the catabolic capabilities of several environmental Pseudomonads. For this we used four well studied model organisms: *Pseudomonas fluorescens* SBW25^30^, *Pseudomonas putida* KT2440^31^, *Pseudomonas protegens* Pf-5^32^, and *Pseudomonas stutzeri* DSM4166^33^. *Sorghum bicolor* is well established as a producer of multiple BNIs, including methyl 3-(4-hydroxyphenyl) propionate (MHPP) – the first PPME found to inhibit biological nitrification^14^. Thus, we also evaluated three recently isolated sorghum endophytes. We utilized the Genome Taxonomy Database toolkit to assign taxonomic classifications for each endophyte ^34^. Based upon these results we classified strain TBS10 as a *Pseudomonas frederiksbergensis* isolate. TBS28 and TBS49 lacked sufficient similarity with existing species to assign a classification at the species level. TBS49 was related to the *P. putida* group, and we refer to it as *Pseudomonas sp.* TBS49. Based upon a combination of its ease of manipulation, diverse catabolic functions (unpublished data), and exceptional genetic tractability we assigned TBS28 the provisional species name *Pseudomonas facilor*.

Each Pseudomonad was screened for their ability to utilize phenylpropanoids. First, each strain was evaluated for use of either glucose or *trans*-*p-*coumaric acid as a sole carbon source (**Fig. 2a**). The phenylpropanoid *p*-coumaric acid is commonly found in plant root exudates and is structurally similar to many PPMEs. All strains grew robustly with glucose as a sole carbon source, but only KT2440, SBW25, TBS10, and TBS28 were able to use *p-*coumaric acid as a sole carbon source. Of the seven, only these four bacteria harbor genes encoding an established pathway for *p-*coumaric acid catabolism (**Supplementary Fig. S1**), and thus these results are unsurprising.

**Figure 2.**
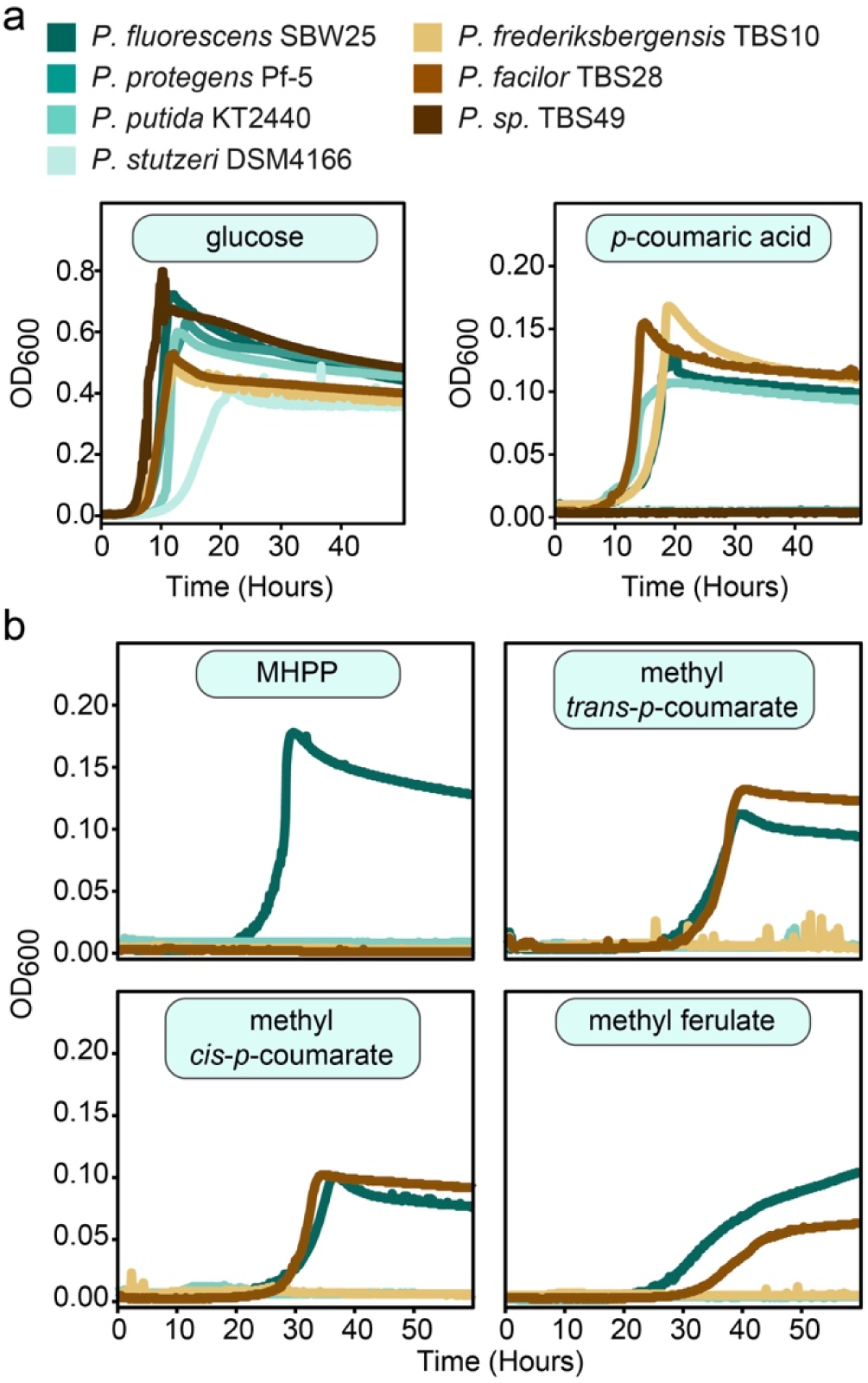
A subset of environmental Pseudomonas isolates can utilize phenylpropanoid methyl esters as carbon sources. Microtiter plate cultivation assays comparing growth of environmental Pseudomonads using (a) two control carbon sources and (b) four phenylpropanoid methyl esters. Experiments in panel b only include the four Pseudomonads that grew with p-coumaric acid in panel a. Assays were performed with MME medium containing either 10 mM (glucose) or 2.5 mM (all others) of the indicated carbon sources. Each panel contains a single representative curve from one of three biological replicates.

The four strains that utilized *p-*coumaric acid were further evaluated for their ability to use four distinct PPMEs (**Fig. 2b**) and phloretic acid (**Supplementary Fig. S2**) as sole carbon sources. The four PPMEs that we used were methyl *trans*-*p*-coumarate, methyl *cis*-*p*-coumarate, methyl ferulate, and MHPP. Phloretic acid is a potential catabolism intermediate of MHPP. SBW25 was able to utilize all four PPMEs as carbon sources, TBS28 used all except MHPP, and the remaining two Pseudomonads were unable to utilize any of the PPMEs. While unable to utilize any PPMEs, KT2440 was able to utilize phloretic acid. Ultimately, SBW25 was chosen for further evaluation due to its ability to use all tested PPMEs.

### Mutant fitness assays identify genes important for robust growth when using PPME as carbon sources

To identify genes involved in catabolism of PPMEs, we utilized high-throughput random barcode transposon site sequencing (RB-TnSeq). RB-TnSeq a powerful tool for identifying genes whose functionality is important during a variety of cultivation conditions. By measuring the change in relative abundance of barcoded transposon mutants within a population one can measure the impact of disrupting gene expression upon cellular fitness. Changes in relative abundance for mutants harboring disruptions for each gene are represented as a fitness value. Generally, negative fitness values indicate the importance of the encoded protein for growth in the test condition and positive fitness scores indicate that abolishing expression of the encoded protein results in faster or more robust growth in the test condition. For this work, we generated a barcoded transposon mutant library based upon *P. fluorescens* JE4621, which is a SBW25-derivative containing a genome integrated poly-*attB* cassette that enables use of serine recombinase-assisted genome engineering (SAGE) ^35^.

The mutant library was profiled for fitness during cultivations with an array of carbon sources. These assays used multiple phenylpropanoids, 4-hydroxybenzoic acid (an intermediate in degradation of *p-*coumaric acid), or glucose (an unrelated carbon source control) as sole carbon sources. A fitness value of −1.8 is typically used as a cutoff to predict whether a gene is important for growth in the test condition (e.g., important for substrate catabolism or stress tolerance) ^36,37^. For experiments in each condition, we identified between 82 (glucose) and 271 (MHPP) genes with fitness values lower than this cutoff (**Supplementary File F1**). Of note, when MHPP was used as a carbon source, we observed far more genes (271 genes) with low fitness than when using any of the other carbon sources (82 to 132 genes). Alternately, high positive fitness values can be indicative of improved growth relative to the bulk population, but are typically uncommon. Accordingly, only a few (0 to 18) genes were found to have a substantial positive fitness value (>1.8) in experiments using most of the carbon sources. However, when using MHPP as a carbon source, we identified 52 genes with this same cutoff.

RB-TnSeq typically allows one to identify a small collection of genes that are directly involved in catabolism of specific carbon sources by comparing fitness values obtained during growth with many different related and unrelated carbon sources. However, as we observe here this type of analysis is not always sufficient. Many genes have substantial negative fitness values across all or most conditions, and thus are likely involved in general cellular processes that are important regardless of carbon source (e.g., biosynthesis of amino acids in minimal medium). These genes represent a ‘background’ signal can be removed from consideration. Indeed, most genes with negative fitness during growth on glucose, also have similar negative fitness with all evaluated carbon sources, and those with negative fitness values unique to glucose were highly enriched for those known to be involved in glucose catabolism. However, this ‘background subtraction’ approach fails when far more genes have substantial fitness scores under one condition (e.g., MHPP) than in other conditions. Even after removing all genes with common fitness values from consideration, there were too many genes to reasonably evaluate on an individual basis for their potential roles in MHPP catabolism. Overall, the exceptionally high number of genes with either substantial positive or negative fitness values during experiments using MHPP suggests that in addition to serving as a carbon source, it may directly or indirectly (e.g., catabolic intermediates such as formaldehyde may be toxic) impact many aspects of cellular physiology. Thus, we sought to reduce the number of genes to evaluate for involvement in PPME catabolism by comparing genes identified by RB-TnSeq with those identified by an orthogonal approach.

### Differential RNAseq reduces the search space for genes involved in catabolic pathways

Differential gene expression analysis (RNAseq) and Random Barcode Transposon-site Sequencing (RB-TnSeq)^37^ are two powerful, orthogonal methods for identifying genes involved in novel metabolic pathways^37–42^ that are very synergistic and enabled PPME pathway discovery that would have been challenging if either was use alone. Despite the fact that each method covers the weaknesses of the other, to our knowledge the two methods have not been previously used in concert to identify catabolic pathways. For example, genes involved in catabolism of related, but distinct compounds may have their expression induced by the compound, or genes involved in catabolism of the compounds may not be differentially expressed (e.g., transcription factors). Alternatively, as RB-TnSeq only evaluates a single gene per cell, it works best when there is no functional redundancy in gene function. Furthermore, phenomena such as metabolite cross-feeding between mutants in the pooled population can result in fitness values that are not representative of the impact of gene deletion in an axenic culture.

We performed RNAseq for SBW25 cultures grown using either glucose or MHPP as carbon sources. We identified 43 genes that were upregulated by at least 8-fold in MHPP cultures versus glucose cultures (**Supplementary File F2, Supplementary Fig. S3**). Genes in the *ech* gene cluster (PFLU3296-3303), which includes genes encoding enzymes involved in *p-*coumaric acid catabolism, are upregulated by MHPP (**Supplementary Fig. S1, Table 1**). We also identified another large cluster of genes (PFLU3304-3311) that is highly upregulated by MHPP (**Table 1**). While most of the *ech* gene cluster is highly conserved between the four *p-*coumarate catabolizing Pseudomonads at both the level of protein sequence (**Supplementary Table S1**) and gene synteny (**Supplementary Fig. S4a**), the additional gene cluster is only partially represented in the sorghum isolates and completely absent in KT2440 (**Supplementary Fig. S4a**). Genes in this cluster are predicted to encode two putative oxidoreductases, a serine hydrolase, several transcription factors, multiple transporters, and two hypothetical proteins.

**Table 1.** Differential expression values and fitness values for genes in the p-coumaric acid and adjacent gene cluster.

| Locus tag | log2 Fold Change (MHPP vs. Glucose) | glucose mean fitness | p-coumaric acid mean fitness | phloretic acid mean fitness | 4-HB mean fitness | ferulic acid mean fitness | MHPP mean fitness |
| --- | --- | --- | --- | --- | --- | --- | --- |
| PFLU3296 | 7.47 | -0.091 | -0.150 | -4.475 | -0.206 | -0.157 | -5.092 |
| PFLU3297 | 7.78 | -0.058 | 0.036 | -2.962 | -0.064 | -0.181 | -4.196 |
| PFLU3298 | 8.09 | 0.130 | -3.411 | -4.670 | 0.323 | -2.061 | -5.189 |
| PFLU3299 | 8.40 | 0.015 | -3.936 | -5.676 | 0.252 | -2.815 | -5.830 |
| PFLU3300 | 8.26 | -0.097 | -3.953 | -4.491 | -0.084 | -3.875 | -5.190 |
| PFLU3301 | 3.17 | --- | --- | --- | --- | --- | --- |
| PFLU3302 | 8.31 | -0.009 | 3.421 | 0.155 | 0.091 | 5.020 | -0.749 |
| PFLU3303 | 7.73 | -0.022 | 0.013 | -0.092 | -0.013 | -0.029 | -1.194 |
| PFLU3304 | 6.52 | 0.054 | -0.153 | -0.152 | -0.064 | 0.007 | 0.216 |
| PFLU3305 | 5.66 | -0.085 | -0.154 | -0.150 | 0.061 | -0.178 | 0.282 |
| PFLU3306 | 6.99 | 0.118 | -0.095 | -0.060 | 0.213 | -0.018 | 0.972 |
| PFLU3307 | 2.52 | 0.165 | 0.078 | 0.054 | 0.460 | 0.129 | 0.933 |
| PFLU3308 | 2.46 | 0.025 | -0.113 | -0.034 | -0.010 | -0.002 | -4.288 |
| PFLU3309 | 7.43 | -0.060 | -0.051 | -0.030 | -0.120 | -0.110 | 0.642 |
| PFLU3310 | 8.14 | 0.076 | 0.066 | 0.193 | 0.000 | -0.083 | 0.522 |
| PFLU3311 | 7.69 | 0.012 | 0.047 | -0.039 | 0.006 | 0.031 | 2.722 |

\*No fitness data available

### Combining RB-TnSeq and RNAseq reduces the search space for novel metabolic enzymes

We were able to reduce the number of genes likely involved in MHPP catabolism from 323 (RB-TnSeq) and 43 (RNAseq) genes down to only 4 genes whose encoded proteins both lacked functional characterization in other organisms and had generic functional annotations (**Fig. 3A**). For this, we directly compared differential expression data (MHPP vs. glucose) with fitness values obtained with each carbon source (**Fig 3**, **Supplementary Fig. S5**). Unsurprisingly, we found that genes encoding enzymes the *p-* coumaric acid catabolic pathway were both upregulated by MHPP and had negative fitness values for cultures grown using any tested phenylpropanoid carbon source (see white dots in lower right sextants of plots with fitness values for phenylpropanoids). Of note, none of the aromatic catabolism genes have substantial fitness values when glucose is the carbon source (**Fig. 3D**).

**Fig. 3.**
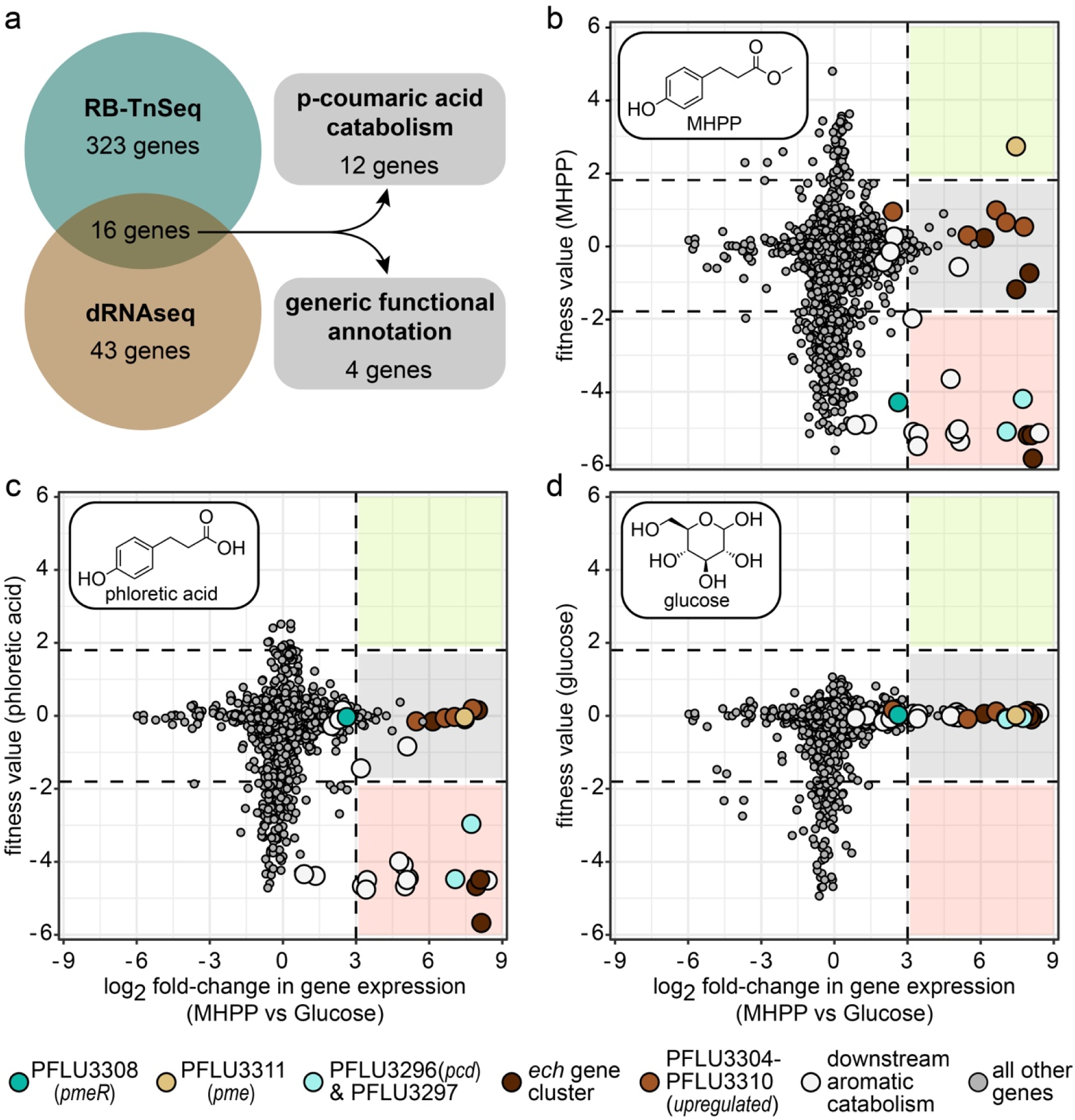
Comparing fitness values with differential gene expression reduces search space for novel pathway genes. (a) Venn diagram showing overlap between genes with substantial fitness values when SBW25 is grown using MHPP and genes that are more highly expressed when SBW25 is cultivated in MHPP versus glucose carbon sources. (b-d) Plots comparing differential expression (x-axis) versus mean RB-TnSeq fitness values on the y-axis. RB-TnSeq values displayed are from cultures grown with either (b) MHPP, (c) phloretic acid, or (d) glucose. Positive and negative differential expression values indicate higher expression during growth using MHPP and glucose as carbon sources, respectively. Dots indicate genes encoding the putative PPME-sensitive pmeR transcription factor (dark teal), phenylpropanoid methyl esterase (light brown), putative phloretoyl-CoA dehydrogenase and putative β-ketothiolase (light teal), other genes in the ech gene cluster (dark brown), other genes in the MHPP-upregulated gene cluster (medium brown), downstream aromatic catabolic pathway gene clusters (white), and all other genes (dark gray). Values represent the mean of 4 RB-TnSeq or 4 differential expression biological replicates. Genes lacking fitness or differential expression values are not displayed.

To identify genes specifically involved in PPME catabolism we narrowed our list of candidate genes to those which satisfy two conditions: (1) they must have substantial fitness values in experiments using PPME or phloretic acid carbon sources, but not in experiments using other aromatic carbon sources; (2) their expression must be substantially higher in cultures grown using MHPP than those grown using glucose (see non-white dots in the top right and bottom right sextants of charts in **Fig. 3B-C** and **Supplementary Fig. S5**). While similar fitness values were observed all aromatic carbon sources for most MHPP-upregulated genes, we observed PPME or phloretic acid-specific fitness values for four MHPP-upregulated genes.

First, the fitness of PFLU3296 and PFLU3297 mutants, the last two genes in the *ech* operon in SBW25, was dependent upon whether MHPP or its potential metabolic intermediate phlore tic acid were the carbon source. The function of the enzymes encoded by the first three genes in the *ech* operon (PFLU3300 to PFLU3298) and their roles in *p*-coumaric acid catabolism are well established in *P. putida* KT2440 and well supported by the combination of both our RB-TnSeq (**Supplementary File F1**) and RNAseq (**Supplementary File F2**) data in SBW25. However, the functions of enzymes encoded by the remaining two genes have yet to be established. PFLU3296 and PFLU3297 encode putative acyl-CoA dehydrogenases and β-ketothiolases, respectively. Substantial negative fitness values were observed for each gene when MHPP or phloretic acid were used as carbon sources (**Fig. 3** – see light teal dots), but fitness values were near zero when using other phenylpropanoid carbon sources. This suggests that these enzymes participate in catabolism of phenylpropanoids with saturated propyl groups such as MHPP and phloretic acid.

To validate whether these two genes participate in MHPP and phloretic acid catabolism we generated targeted gene deletions in *P. fluorescens* JE4621 ^43^ and evaluated the ability of the resulting strains to use each carbon source (**Fig. 4B**). While deletion of PFLU3296 abolished growth with MHPP and phloretic acid, deletion of PFLU3297 had no impact. We attempted complementation of the PFLU3296 deletion by reintroducing a copy of PFLU3296 under control of a constitutive promoter into the chromosome of the deletion strain with SAGE^43^. The complemented strain was able to utilize both MHPP and phloretic acid, demonstrating that lack of PFLU3296 expression (**Supplementary Fig. S6**), rather than a polar effect was the cause of the phenotype observed in the PFLU3296 deletion strain. As the PFLU3296 deletion clearly induces a growth phenotype, we suspect that the low fitness score of PFLU3297 in the RB-TnSeq data may be a consequence of a polar effect that influences PFLU3296 expression. However, this remains unconfirmed.

**Fig. 4.**
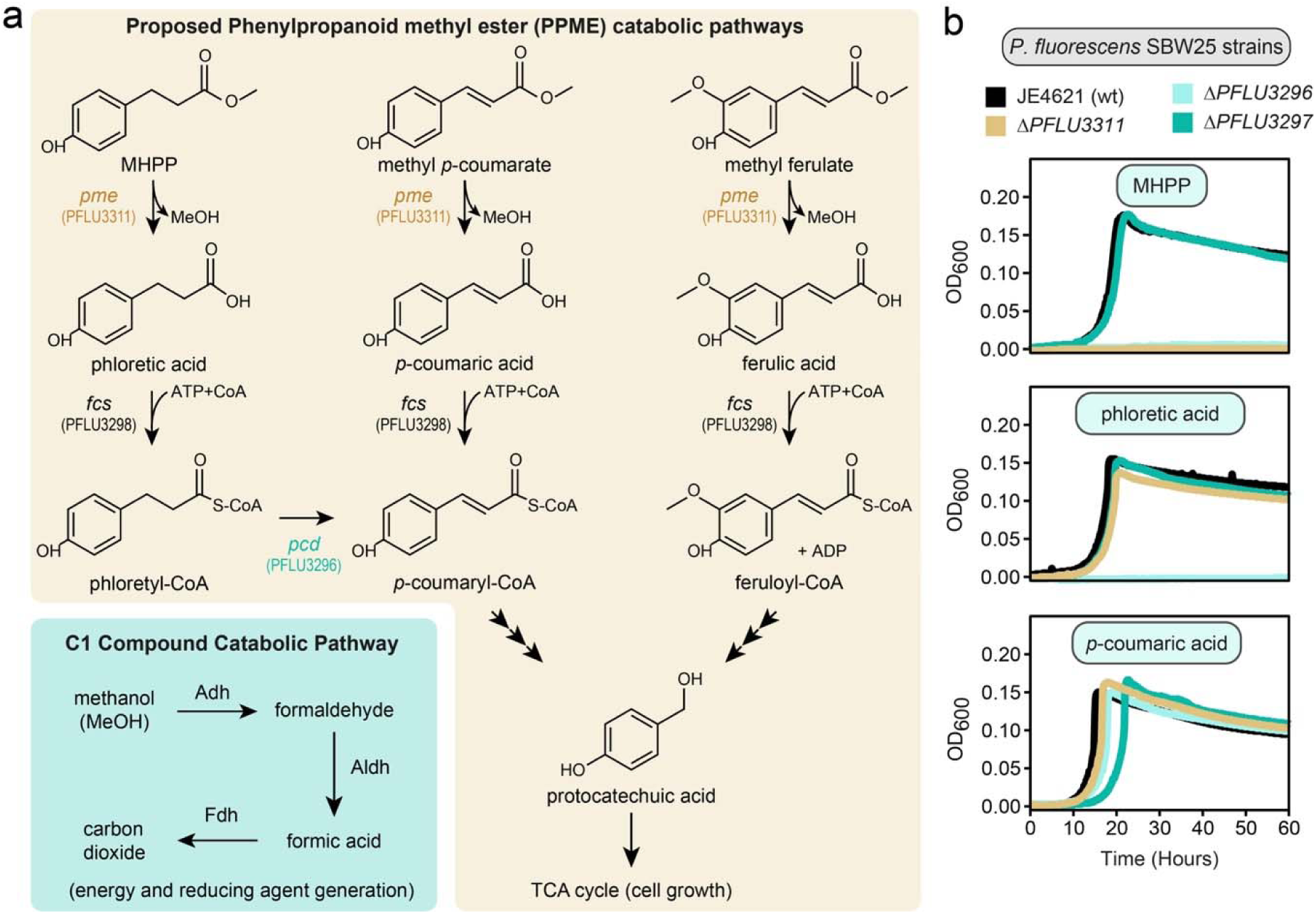
Genetic dissection of a proposed pathway for PPME catabolism. (a) Proposed pathways for catabolism of several different PPMEs. Brown highlighted pathways follow catabolism of the phenylpropanoid moeity following an initial de-esterification step, and teal highlighted pathway is a standard C1 metabolic pathway that follows the MeOH moeity released upon de-esterification. (b) Microtiter plate cultivation comparing growth of P. fluorescens SBW25-derived targeted gene deletion mutants. Assays were performed with MME medium containing 2.5 mM each of the indicated carbon sources. Each panel contains a single representative curve from one of three biological replicates.

Second, the fitness of PFLU3308 and PFLU3311 mutants was highly dependent upon whether MHPP was the carbon source. Expression of PLFU3308, which encodes a transcription factor, does not quite reach our 8-fold differential gene expression threshold. However, PFLU3308 has a negative fitness with MHPP that is not observed with any other tested carbon source (**Fig. 3B, Supplementary Fig. S5** – see dark teal dot). This suggests a role in regulating expression of genes required for PPME catabolism. Inactivation of a critical enzyme in a catabolic pathway typically abolishes use of the carbon source. In heterogenous RB-TnSeq mutant libraries, such mutants become reliant upon metabolites secreted by other strains for growth, and thus generally have low fitness values. However, no other genes are both upregulated by MHPP and have negative fitness values associated with only MHPP alone. One possible explanation for this lack is that multiple redundant enzymes may convert PPMEs into *p-*coumaric acid pathway intermediates, and thus no individual enzyme is essential for this conversion. Another is that enzyme(s) performing this step are not differentially expressed – which seems unlikely, as such an enzyme would not likely have a role in growth with glucose as a carbon source.

PFLU3311, which encodes a putative serine hydrolase, is both highly upregulated by MHPP, and has a highly *positive* fitness value when MHPP is the carbon source (**Fig. 3B, Supplementary Fig. S5** – see light brown dot in top right sextants). So, a third possible explanation for the lack of an enzyme-encoding gene with a negative fitness value unique to MHPP is that the first step of catabolism may release a toxic intermediate that reduces fitness. If PFLU3311 is responsible for hydrolyzing the ester bond in PPMEs, then methanol or other toxic C1 compounds formed by its oxidation (e.g., formaldehyde) will be generated inside the cell and may inhibit growth (**Fig. 4A**). Thus, in a heterogenous population where cross-feeding of other less toxic PPME metabolic intermediates (e.g., *p-*coumarate or 4-hydroxybenzoate) can occur, a mutant that does not produce these C1 compound may have a growth advantage (indicated by positive fitness value). However, if the enzyme encoded by PFLU3311 does perform the first step in the pathway, one would expect growth of a PLFU3311 mutant on PPMEs to be abolished in an axenic culture. Indeed, when PFLU3311 is deleted from JE4621, the resulting mutant is unable to grow with PPMEs as a carbon source, but the mutant retains the ability to utilize phloretic acid (**Fig. 4B**). Finally, we observed that reintroduction of PFLU3311 into the deletion mutant with SAGE restored the ability to use PPME carbon sources – thus ruling out polar effects as the cause of the mutant growth phenotypes (**Supplementary Fig. S6**).

Depending upon the phenylpropanoid carbon source, we observed fitness values for PFLU3302 that range from highly positive to mildly negative. PFLU3302 encodes a putative porin, and its disruption by transposons results in highly positive fitness values when using ferulic acid or *p-*coumaric acid carbon sources as carbon sources. However, when using other phenylpropanoids the fitness was either neutral or mildly negative. While the high fitness value indicates that the PFLU3302 mutant had a growth advantage over other mutants in the RB-TnSeq library, we observed no change in growth phenotypes in a targeted mutant when using *p-*coumaric acid, phloretic acid, or MHPP as carbon sources carbon sources (**Supplementary Fig. S7**). This suggests that disruption of PFLU3302 expression only provides a conditional benefit. We also observed moderate negative fitness values for PFLU3303 during growth using MHPP, but targeted deletion had no impact on growth with tested phenylpropanoids (**Supplementary Fig. S7**). Together these data suggested there is likely a significant amount of redundancy in phenylpropanoid transport.

In addition to genes encoding proteins that may be directly involved in catabolism of PPMEs, we also observed several putative stress response genes whose disruption was reduced fitness with MHPP, but not other aromatic carbon sources. PFLU0397-0399, which encode orthologs of the stress activated HslUV protease from *Escherichia coli* and a downstream gene of unknown function, had strong negative fitness values when MHPP is the carbon sources. Orthologs of HptX protease (PFLU1730), HptG molecular chaperone (PFLU1830), and SoxR superoxide stress-sensitive transcription factor (PFLU2053) all had negative fitness values that were also unique to MHPP. However, none of these genes were highly differentially expressed.

Finally, PFLU3298 (*fcs*) and PFLU3300 (*ech*) encode the enzymes responsible for the first two steps in the *p-*coumaric acid catabolism pathway (**Supplementary Fig. S1**). As predicted by our RB-TnSeq results and previous work in *P. putida* ^36^, disruption of either of these two genes with transposons prevented use of any phenylpropanoid carbon sources (**Supplementary File F1**). Thus, conclusively demonstrating that PPMEs are catabolized through the established phenylpropanoid catabolism pathway.

### PFLU3311 encodes a representative of a novel phenylpropanoid methyl esterase enzyme family

Given its importance in PPME catabolism, we sought to determine the biochemical function of the putative serine hydrolase enzyme encoded by PFLU3311. Serine hydrolases are one of the largest classes of enzymes^44,45^ with functions spanning from proteases ^46^ to hydrolysis of the ester linkages in polyethylene terephthalate by PETase^47^. The protein encoded by PFLU3311 has low sequence homology with any well characterized enzymes (Supplementary Fig. S8 – protein similarity tree), including a functionally similar feruloyl esterase family that is thought to be responsible for breaking the phenylpropanoid-sugar ester linkages found between lignin and hemicellulose ^48^. The most similar proteins with an experimentally validated function are two 6-aminohexanoate dimer hydrolases that cleave amide bonds in linear nylon oligomers^49,50^. However, the sequence similarity with even these proteins was below 35% identity with 88% query coverage when using BLASTp (see **Supplementary Table S2** for protein sequences) ^51^. This suggests that PFLU3311 likely represents a novel family of enzymes. Above, we posited that the first step in catabolism of PPMEs may be hydrolysis of the ester bond, resulting in generation of their cognate PP carboxylic acids and methanol. Thus, we evaluated the ability of the PFLU3311 serine hydrolase to hydrolyze PPMEs into their cognate carboxylic acids with *in vitro* biochemical activity assays. We expressed a codon-optimized version of PFLU3311 containing a C-terminal 6x-His-tag in *Escherichia coli* BL21(DE3) pLysS, purified the protein by immobilized metal affinity chromatography, and evaluated the ability of the resulting purified protein to hydrolyze four PPMEs by LC-MS (**Fig. 5**). Unlike feruloyl esterases, which show poor, but measurable, activity with PPMEs ^48^, the enzyme encoded by PFLU3311 completely converts each PPME into their cognate carboxylic acid. This conclusively demonstrates that PFLU3311 encodes a phenylpropanoid methyl esterase.

**Fig. 5.**
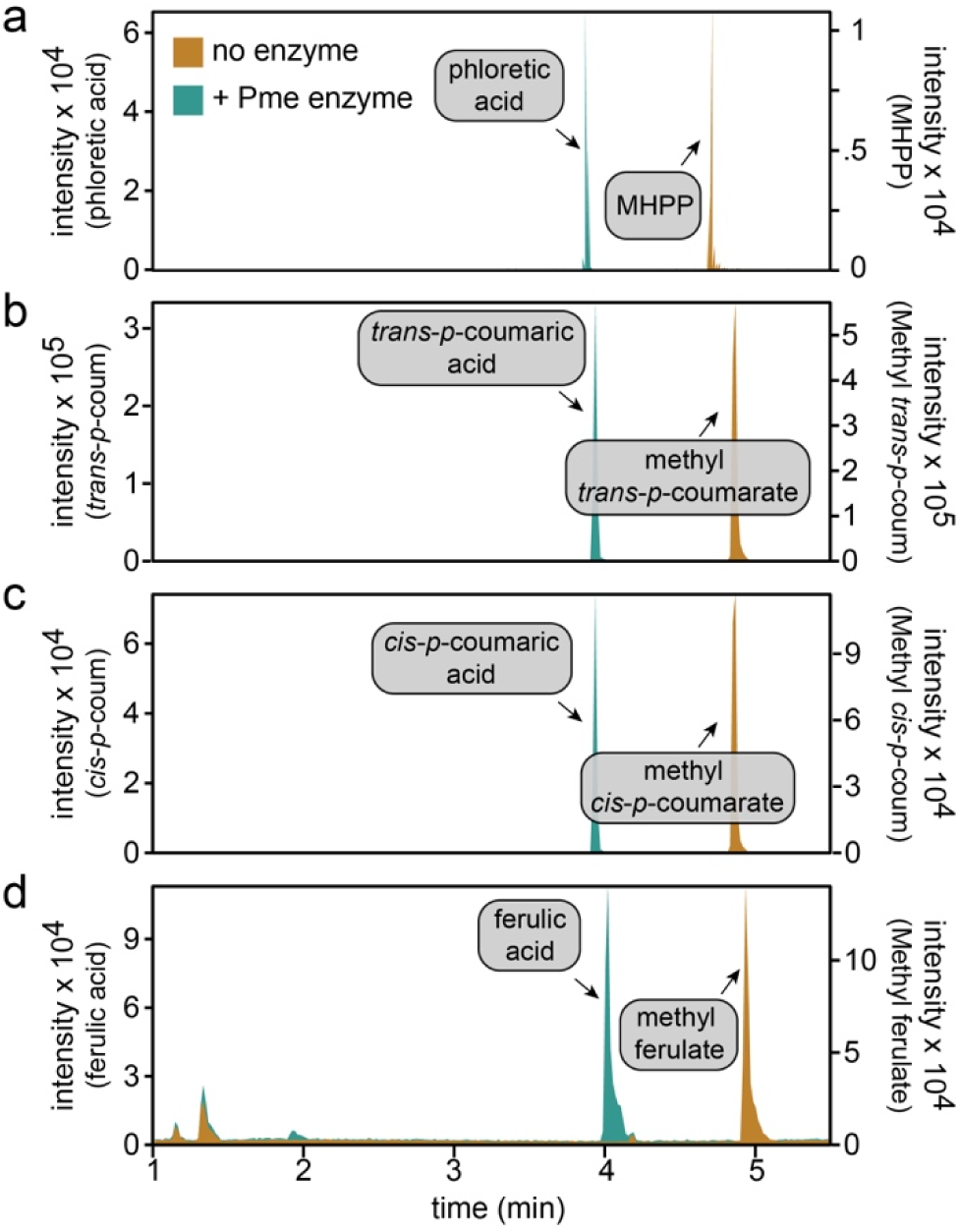
PFLU3311 encodes a phenylpropanoid methyl esterase. LC-MS chromatograms of products in enzyme assay samples containing (a) MHPP, (b) methyl-trans-p-coumarate, (c) methyl-cis-p-coumarate, and (d) methyl ferulate. Chromatograms of samples with and without the protein encoded by PFLU3311 are colored teal and brown, respectively. For each compound other than methyl ferulate, there is a single predominant ion and thus for panels a-c the intensity is represents base peak intensity. Methyl ferulate had two prominent peaks, and thus intensity represents the total ion count. Each displayed chromatogram is a single representative of 3 replicate enzyme activity assays.

### A putative serine hydrolase and acyl-CoA dehydrogenase are critical for catabolism of PPMEs

Based upon these results we propose a pathway for PPME catabolism (**Fig. 4A**). First, phenylpropanoid methyl esterase, encoded by the *pme* gene (PFLU3311), hydrolyzes the PPME, releasing a carboxylic acid and methanol. The methanol moiety is likely catabolized via existing C1 oxidation pathways (**Fig. 4A** – teal highlight). The aromatic acids produced by this reaction are ligated with CoA by feruloyl-CoA synthetase and processed through parallel pathways (**Fig. 4A** – brown highlight). Ferulic acid and *p-*coumaric acid are subsequently metabolized into TCA cycle intermediates via established pathways. In our proposed pathway phloretoyl-CoA, the product of CoA ligation to phloretic acid, is subsequently oxidized by <u>p</u>hloretoyl-<u>C</u>oA <u>d</u>ehydrogenase, which is the product of *pcd* (PFLU3296), into *p-*coumaroyl-CoA which is then catabolized by the same pathway used for *p-*coumaric acid catabolism.

### Heterologous expression of PPME catabolic pathways to elucidate gene function

As an orthogonal approach to support the pathway proposed above, we sought to engineer *Pseudomonads* to use PPMEs that they do not natively catabolize as carbon sources. *P. putida* KT2440, *P. facilor* TBS28, and *P. frederiksbergensis* TBS10 each contain a subset of the genes we identified as being involved in PPME catabolism (**Supplementary Fig. S4**), and thus transfer of PFLU3311 and PFLU3296 into these organisms will serve as an orthogonal evaluation of their function in PPME catabolism. For this, we used Bxb1 recombinase to transfer constitutively expressed copies of PFLU3311, PFLU3296, or both into the chromosomes of KT2440, TBS28, and TBS10 variants, respectively (**Fig. 6**)^43,52^. The *ech* operon in KT2440 has the same gene arrangement and composition as SBW25, and it contains orthologs of both PFLU3296 and PFLU3297 (**Supplementary Fig. S4**). The *ech* operon in TBS28 and TBS10 contain the first three genes found in the SBW25 *ech* operon, but they lack orthologs of both PFLU3296 and PFLU3297. Unlike KT2440 and TBS10, the TBS28 genome harbors an ortholog of PFLU3311. The *pme* gene in SBW25 is in a gene cluster adjacent to the *ech* operon – which aided in inferring a potential role in PPME catabolism. However, the TBS28 *pme* ortholog is distal from the *ech* operon, is separated from orthologs of most other genes in the SBW25 gene cluster and has no ortholog of the transcription factor encoded by PFLU3308 – which appears to be essential for regulating genes essential for PPME catabolism in SBW25. Thus, discovery of *pme* in TBS28 would have likely proven even more challenging than SBW25. Together, these genetic features likely explain why (1) KT2440 catabolizes phloretic acid, but not PPMEs; (2) TBS28 catabolizes all PPMEs other than MHPP and cannot catabolize phloretic acid; (3) TBS10 catabolizes *p-*coumaric acid, but none of the other compounds.

**Fig. 6.**
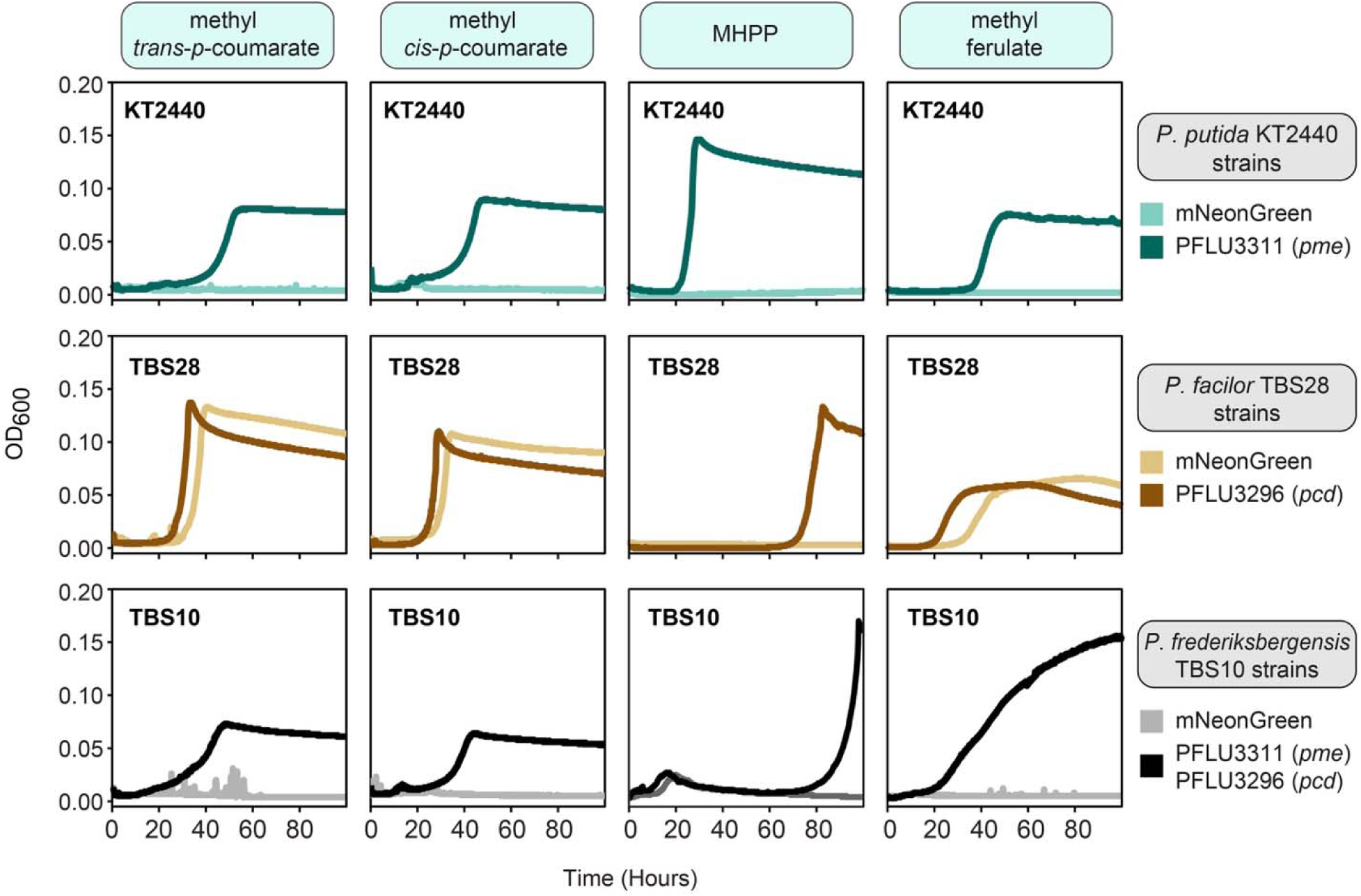
Transfer of pme and pcd genes enables heterologous hosts to use PPMEs as carbon sources. Microtiter plate cultivation assays comparing growth of Pseudomonads expressing either the fluorescent protein mNeonGreen or the Pme/Pcd enzymes from SBW25. Assays were performed with MME medium containing 2.5 mM of the indicated carbon source. Each panel contains a single representative curve from one of three biological replicates.

Indeed, we found that expression of the SBW25 genes PFLU3311, PFLU3296, or the combination of the two, but not an mNeonGreen fluorescent reporter negative control enabled each strain to utilize PPME carbon sources (**Fig. 6**). However, TBS10, and to a lesser extent TBS28, catabolized MHPP slowly when expressing *pme* and *pcd*. Heterologous expression of a transporter from SBW25 was not required for growth in these strains. However, the slow growth in TBS28 and TBS10 may be indicative of poor activation of downstream pathways (e.g., poor expression of Fcs) or inefficient transport – which may be due to poor expression or low substrate affinityTaken together, this solidifies the finding that the enzymes encoded by these two genes are critical for catabolism of PPMEs and demonstrates a relatively low barrier for horizontal transfer of PPME catabolic capability.

## Discussion

Enhancing biological nitrification inhibition has been proposed as a tool to reduce agricultural nitrogen pollution ^13^, but many of the underlying mechanisms remain poorly understood. It is currently difficult to develop assays to probe mechanisms other than simple enzyme inhibition because the rhizosphere is such a complex system. The process by which BNI compounds are removed from soil is also not understood. A deeper understanding of this process will be essential for both developing a tractable model system for studying BNI mechanisms and engineering rhizobacteria to produce BNI compounds. Here, we identify a previously unknown catabolic pathway that is used by *P. fluorescens* SBW25 to catabolize the PPME class of BNI compounds. Wild-type SBW25, its mutants, or other rhizosphere bacteria engineered with BNI catabolism pathways can be used to both study how temporal mineralization of BNI compounds affects (de)nitrification and evaluate whether microbe-microbe interactions between AOA/AOB and BNI-consuming microbes affects nitrification. Finally, this work provides a pathway for identifying organisms which catabolize other BNI compounds (e.g., sorgoleone) and elucidating their catabolic pathways, which will be essential to understanding how BNI works on a broader scale.

In this study, we used a two-pronged approach combining RNA-Seq with RB-TnSeq for pathway discovery. Both RNA-Seq and RB-TnSeq have been used independently to elucidate catabolic pathways in bacteria^37–42^. In our study, almost 5% of the genes in SBW25 were greater than 4-fold differentially expressed between cultures grown using MHPP and glucose carbon sources. With RB-TnSeq, a similar amount of genes had substantial fitness values. Even after disregarding genes that are likely involved in general metabolic processes the remaining number of genes to evaluate too high to reasonably evaluate by targeted gene deletion – the standard method for validating gene involvement in novel pathways. With a combined approach, we were able to substantially narrow the list to testable number of genes and overcome the technical limitations of each approach. Catabolic pathways for aromatic compounds tend to converge into small group of established pathways^53,54^, such as the phenylpropanoid catabolic pathway used for *p-*coumaric acid. So, we expected that MHPP may be funneled into one of these established pathways. However, relationships with established catabolic pathways are often unavailable when evaluating catabolic pathways for new compounds, and many compounds of recent interest (e.g., sorgoleone, lignin oligomers, polyurethanes, *etc.*) are substantially more complex than PPMEs. As such, we expect that combining these two orthogonal high-throughput ‘omics methods for catabolic pathway discovery will accelerate characterization of these complex pathways.

Pme in *P. fluorescens* and *P. facilor* represent the first characterized enzymes in a family of novel methyl esterases and serve as the key enzyme for catabolism of PPMEs. While the catabolic pathway for the aromatic component of PPMEs is now elucidated, our results did not strongly suggest a role for specific existing C1 pathways in catabolism of the methanol moiety produced by de-esterification of PPMEs. In Pseudomonads, pyrroloquinone (PQQ) dependent methanol dehydrogenases that are localized to the periplasm are the primary enzymes responsible for catabolism of short chain *n*-alcohols^55,56^. However, we did not see upregulation of these enzymes or fitness defects following disruption of these enzymes in our data. This may be a consequence of functional redundancy, as has been observed previously in *P. putida*^55^, or suggest that other alcohol dehydrogenase(s) function on intracellular methanol. Thus, it remains unclear how intracellular methanol is catabolized in these organisms. It is possible that basal levels of multiple alcohol and aldehyde dehydrogenases provide sufficient expression of functionally redundant dehydrogenase enzymes. Without dedicated enzymes the slow mineralization of methanol to CO_2_ may explain the apparent toxicity associated with Pme expression in the presence of MHPP. Nonetheless, this remains an open question for future research to address.

*P. fluorescens* SBW25 was isolated from a beet rhizosphere, and thus it was somewhat surprising that it was able to utilize a BNI compound that is only currently known to be exuded by sorghum roots ^14^. Interestingly, none of the sorghum endosphere isolates were able to utilize MHPP as a carbon source. This may be explained in part by their environmental niches. Timm *et al* found that *P. fluorescens* isolates from the rhizosphere and endosphere of the same poplar plants are enriched for distinct sets of metabolic pathways ^57^. Specifically, pathways for root exudate compounds were substantially less prevalent in endophytes compared to isolates from the same plant rhizosphere. Ultimately, the ability for an isolate from the rhizosphere of a very distinct plant species suggests that exudation of PPMEs by plant roots may be more common than the current literature suggests.

## Materials and Methods

### General culture conditions & media

The strains and plasmids used in this study are listed in **Supplementary Table S3**. Routine cultivation of *Escherichia coli* for plasmid construction and maintenance was performed at 37 °C using LB medium supplemented with 50 µg/mL kanamycin sulfate or 50 µg/mL apramycin sulfate and 15 g/L agar (for solid medium). Cultivations for strain maintenance, competent cell preparations, and starter cultures for all Pseudomonas strains were performed at 30 °C with 200 rpm shaking in LB or SOB (for TBS10).

MME medium (containing 9.1 mM K_2_HPO_4_, 20 mM MOPS, 10 mM NH_4_Cl, 4.3 mM NaCl, 0.41 mM MgSO_4_, 68 µM CaCl_2_, 1x MME trace minerals, pH adjusted to 7.0 with KOH) supplemented with carbon sources as indicated in the text was used for cultures in growth assays, transcriptomics experiments, and fitness assays. 1000x MME trace mineral stock solution contains per liter, 1 mL concentrated HCl, 0.5 g Na_4_EDTA, 2 g FeCl_3_, 0.05 g each H_3_BO_3_, ZnCl_2_, CuCl_2_·2H_2_O, MnCl_2_·4H_2_O, (NH_4_)_2_MoO_4_, CoCl_2_·6H_2_O, NiCl_2_·6H_2_O. Other than glucose, all carbon sources were used at 2.5 mM. Glucose was used at either 10 mM (growth assays and transcriptomics experiments) or 5 mM (fitness assays).

### Plasmid construction

Q5 DNA polymerase (Thermo Scientific) and primers synthesized by Eurofins Genomics were used in all PCR amplifications for plasmid construction. OneTaq® (New England Biolabs - NEB) was used for colony PCR. Plasmids were constructed by Gibson Assembly using NEBuilder® HiFi DNA Assembly Master Mix (NEB) or ligation using T4 DNA ligase (NEB). Plasmids were transformed into NEB 5-alpha F’I^q^ (NEB). Standard chemically competent *Escherichia coli* transformation protocols were used to construct plasmid host strains. Transformants were selected on LB (Miller) agar plates containing 50 µg/mL kanamycin sulfate for selection and incubated at 37 °C. Template DNA was isolated *P. fluorescens* SBW25 or *P. facilor* TBS28 using Zymo Quick gDNA miniprep kit (Zymo Research) or cells were used directly as a template. Zymoclean Gel DNA recovery kit (Zymo Research) was used for all DNA gel purifications. Plasmid DNA was purified from *E. coli* using GeneJet plasmid miniprep kit (ThermoScientific) or ZymoPURE plasmid midiprep kit (Zymo Research). Sequences of all plasmids were confirmed using Sanger sequencing performed by Eurofins Genomics. Plasmids used in this work are listed in **Supplementary Table S3**. Plasmid sequences are in **Supplementary File P1**.

### Pseudomonad strain construction

All genome modifications were performed using either the allelic exchange or by integration of non-replicating plasmids into the chromosome with Bxb1 integrase. Allelic exchange was performed as described previously ^53^ with the exception that pK18sB ^58^ was used instead of pK18mobsacB. Plasmids for expressing PFLU3311, PFLU3296, or PFA28_02292 were integrated into the chromosomes of host strains using Bxb1 integrase. Briefly, Bxb1 integrase unidirectionally recombines DNA between a cognate *attB* and *attP* sequences. Expression plasmids each harbor a Bxb1 *attP* sequence, and variants of *Pseudomonas putida* KT2440 (JE90), *Pseudomonas fluorescens* SBW25 (JE4621), *Pseudomonas fredericksbergensis* TBS10 (JE5041), and *Psuedomonas facilor* TBS28 (RS175) have each been engineered to harbor a cognate Bxb1 *attB* sequence. Strains JE90, JE4621, and JE5041 were produced previously ^43,52^. Allelic exchange was used to incorporate a DNA cassette that includes multiple *attB* sequences, including Bxb1, downstream of the *ampC* (PFA28_1309) gene in the chromosome of TBS28, generating strain RS175. Insertion of the poly-*attB* cassette into TBS28 was confirmed by both colony PCR and with integration of pGW60 – which includes an mNeonGreen expression cassette – into the Bxb1 *attB* site using SAGE. The resulting strain was named *Pseudomonas facilor* RS175. Deletion of genes in either the *ech* gene cluster or uncharacterized MHPP-upregulated gene cluster was performed using allelic exchange. Gene deletions were confirmed by colony PCR. Mutant strains and the plasmids used for each gene deletion or gene insertion are listed in **Supplementary Table S3**, respectively.

### Plate reader growth assays

6 mL SOB medium (TBS10) or LB (all other strains) was inoculated from glycerol stocks and incubated overnight at 30°C, 200 rpm for precultures cultures. Starter cultures were prepared by inoculating 6 mL MME supplemented with 10 mM glucose from LB precultures (1% inoculum), and incubated at 30°C, 200 rpm until stationary phase was reached (typically overnight) to synchronize cultures, normalize inoculum, and exhaust residual glucose. Growth assays were performed with 600 uL MME supplemented with appropriate carbon sources per well in clear 48-well microtiter plates with an optically clear lid (Greiner Bio-One). Starter culture OD600 values were spot-checked to ensure similar starting inoculum. Assay plates were incubated at 30 °C with 548 rpm with a 2 mM orbital in a Neo2SM (BioTek) or Synergy H1(BioTek) plate reader, with OD_600_ readings taken every 10 minutes. Growth curves displayed in figures were generated using the R-package ggplot2, and are representative curves from 3 biological replicates.

### RB-TnSeq fitness assays

RB-TnSeq was performed according to Wetmore et al. 2015^37^. Briefly, the mutagenized *Pseudomonas fluorescens* JE4621 RB-TnSeq strain libraries were thawed at room temperature and cultured in 25 mL LB media at 30°C with 200 rpm shaking until OD_600_ reached 0.5. The library was adapted to defined medium by inoculating 25 mL MME supplemented with multiple carbon sources (5 mM glucose, 5 mM sodium citrate, 15 mM sodium acetate) with 1% of the LB culture and incubated at 30 °C with shaking until reaching stationary phase. This culture was centrifuged, washed twice with MME, and resuspended to an OD_600_ of 2. Aliquots of washed cells were centrifuged, decanted, and resulting cell pellet stored at −80 °C for later amplicon sequencing and barcode quantification. These aliquots served as T=0 samples for RB-TnSeq. The remaining washed cells were added to 5 mL MME medium containing various carbon sources to reach a starting OD600 of 0.02. These cultures were incubated in 24-well deep well plates at 30 °C and 700 rpm shaking (2 mM orbital) until stationary phase was reached for each carbon source (typically 24-36 hours). Following completion of growth cells were harvested for amplicon sequencing and barcode quantification. Calculated fitness values can be found in Supplemental File F1.

### Transcriptomics experiments and differential expression analysis

*P. fluorescens* SBW25 was cultured at 30 °C in 50 mL MME medium supplemented with either 10 mM glucose or 5 mM MHPP in a 250 mL Erlenmeyer shake flask at 30 °C, 200 rpm shaking and harvested mid-log (OD_600_ = ∼0.2) by centrifugation (∼16,000 x g, 2 minutes, 4 °C). Supernatants were quickly decanted, and cell pellets were frozen rapidly in liquid nitrogen prior to storage at −80 °C. Four samples were prepared for each condition. Cell pellets were submitted to GeneWiz for RNA extraction, 16s rRNA depletion and RNA sequencing.

After Illumina sequencing, RNA-seq reads were mapped to the *P. fluorescens* SBW25 reference genome (NC_012660) using the Geneious for RNA-seq mapping workflow. Read count per annotated gene was calculated for each treatment and replicate, as well as fragment per kilobase million (FPKM), a common normalization technique. We then used the implementation of the DEseq2 R-package that is built into Geneious to calculate differential gene expression. DESeq2 calculates log-fold change in expression and allows comparison between treatments using several ^59^. We had four replicates per treatment. Differential expression data can be found in Supplemental File F2. RNAseq data were deposited in the Gene Expression Omnibus (GEO) under accession number GSE228022.

### Phylogenetic analysis of protein sequences

Protein sequences from functionally validated enzymes and other representative uncharacterized proteins were obtained from GenBank. Protein sequences were aligned through the Aliview (v1.26) alignment viewer and editor^60^ using mafft (v7.453) ^61^ with the –localpair option and a maximum of 1000 iterations. The tree was constructed using FastTree^62^ and visualized using FigTree (http://tree.bio.ed.ac.uk/software/figtree/).

### Recombinant expression and protein purification

Freshly transformed BL21(DE3) *E. coli* colonies harboring pEVF-SBP1-P2, a PFLU3311 expression vector, were used to inoculate starter cultures (5 mL) by scraping a swath of cells from a fresh LB agar plate. Starter cultures were grown overnight in 2x YT medium (no more than 16 hours) in the presence of 50 µg/mL kanamycin sulfate and were used to inoculate expression cultures of ZYM-5052^63^ media (100 mL, 1:100 dilution) containing 50 µg/mL kanamycin sulfate. Expression cultures were grown with constant shaking at 250 rpm at 37°C until reaching an OD_600_ (optical density at 600 nm) of 1.5, wherein they were induced with a final concentration of 0.1% arabinose. Upon induction, the temperature was decreased to 25 °C and cultures incubated for an additional 18 hours before cell harvesting by centrifugation at 5400 x g.

Cell pellets were resuspended in a lysis/wash buffer (50 mM Tris-HCl, 500 mM NaCl, and 5 mM imidazole, pH 7.5) and lysed via sonication. Cell debris was pelleted by centrifugation at 15,000 x g for 25 min at 4°C and clarified cell lysate was recovered. To bind His_6_-tagged protein, clarified cell lysate was incubated with 150 μl of Ni-NTA resin at 4°C for 1 hour with rocking. Resin was collected and extensively washed with 50 resin bed volumes (bv) of lysis buffer. Bound protein was eluted from the TALON resin by incubation with 5 bv of elution buffer (lysis buffer with additional 200 mM imidazole) concentrated to 40 µM and exchanged into 25 mM Tris-HCl, 95 mM NaCl, 15 mM imidazole with a 3-kDa MWCO Vivaspin spin-concentration filter (GE Health Sciences). Protein concentration was determined by absorbance at 280 nm and flash-frozen with liquid nitrogen and stored at −80°C until needed.

### Methyl esterase enzyme assays

Enzyme assays were performed in the following manner. Reactions with and without purified enzyme (500 nM) were prepared in a 500 µL solutions of assay buffer (25 mM Ammonium bicarbonate pH 7.5) containing either MHPP, methyl *trans*-*p-*coumaric acid, methyl *cis-p-*coumaric acid, or methyl ferulic acid (5 mM final) as the substrate. Reactions incubated at 25°C for one hour, and the reaction was terminated by passing the solution through a 3-kDa MWCO Vivaspin spin-concentrator and collecting the filtrate. This reaction was performed in triplicate for each condition (+/− enzyme). The filtrate was then diluted 100-fold in pure water and submitted for LC-MS analysis.

### Liquid Chromatography—Mass Spectrometry Analysis of enzyme assay products

Ten microliters of sample was injected, and analytes were separated on a ZORBAX Extend C18 column (2.1 x 150 mm; 1.8 µm) maintained at 24°C with a flow rate of 300 µL min^-1^at a Waters UPLC system. The mobile phase, comprising water with 0.1% formic acid (A) and acetonitrile with 0.1% formic acid (B) was held at 95% A for the first 1.12 min, ramped to 99.5% B at 6.4 min, and held at 99.5% until 10 min. The mobile phase was held at 95% A for 2 min following data acquisition for post-run conditioning. Mass spectrometry data was acquired with a Waters Xevo G2 QTOF using positive ionization to detect ferulic acid, methyl ferulic acid, and MHPP, and negative ionization for *p*-coumaric acid, methyl coumaric acid, and phloretic acid. Conditions in the spray chamber were as follows: 80°C source temperature; 150°C desolvation temperature; 600 L h^-1^ desolvation gas flow; and 40 V cone voltage. A capillary voltage of 2.5 kV was used, and mass spectra were continuously recorded within a m/z range of 50 – 1200. Compounds were identified by comparison with authentic standards and chromatograms were displayed with base peak intensity (BPI), or total ion current (TIC).

## Acknowledgements

This research was supported in part by the U.S. Department of Energy (DOE), Office of Biological and Environmental Research (BER), as part of BER’s Genomic Science Program (GSP), and is a contribution of the Pacific Northwest National Laboratory (PNNL) Secure Biosystems Design Science Focus Area “Persistence Control of Engineered Functions in Complex Soil Microbiomes”. Additional support was provided by the Laboratory Directed Research & Development program at PNNL. PNNL is a multi-program national laboratory operated by Battelle for the DOE under Contract DE-AC05-76RL01830.

