## Supplementary Materials for "A novel phenylpropanoid methyl esterase enables catabolism of aromatic compounds that inhibit biological nitrification"

### Supplementary Figures

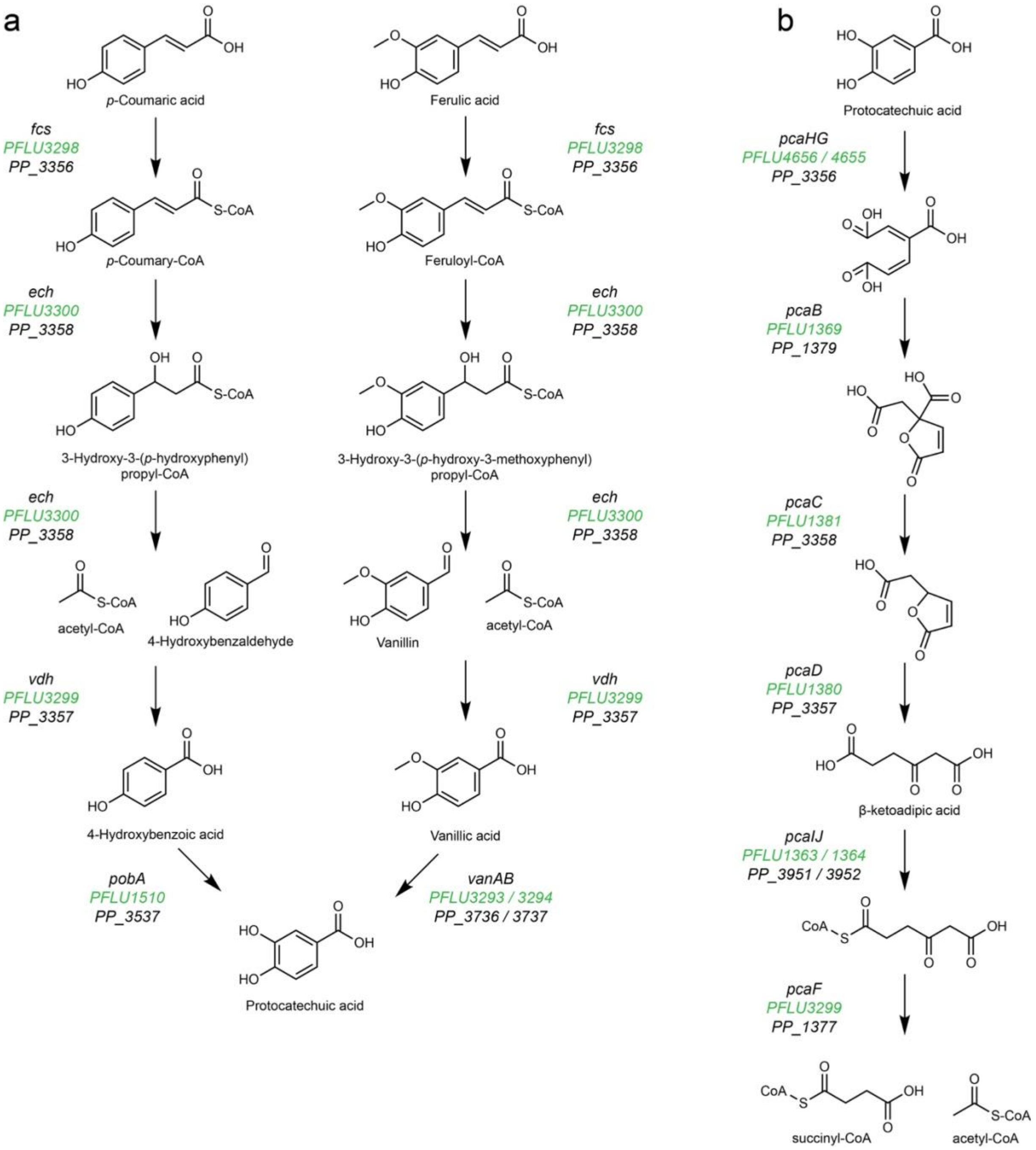


**Supplementary Fig. S1. Catabolic pathways for *p*-coumaric and ferulic acids.** (a) Convergent portion of the catabolic pathways for the two phenylpropanoid carbon sources, which is largely comprised of enzymes encoded in the *ech* gene cluster. (b) Ortho-cleavage pathway for protocatechuic acid, which is also commonly referred to as the B-ketoadipate pathway after its characteristic intermediate. This portion of the pathway funnels the aromatic compounds into the TCA cycle. Names of the genes encoding the enzymes that perform each step are shown in *italics* and locus tags for *P. fluorescens* SBW25 and *P. putida* KT2440 are shown in green and black text, respectively.

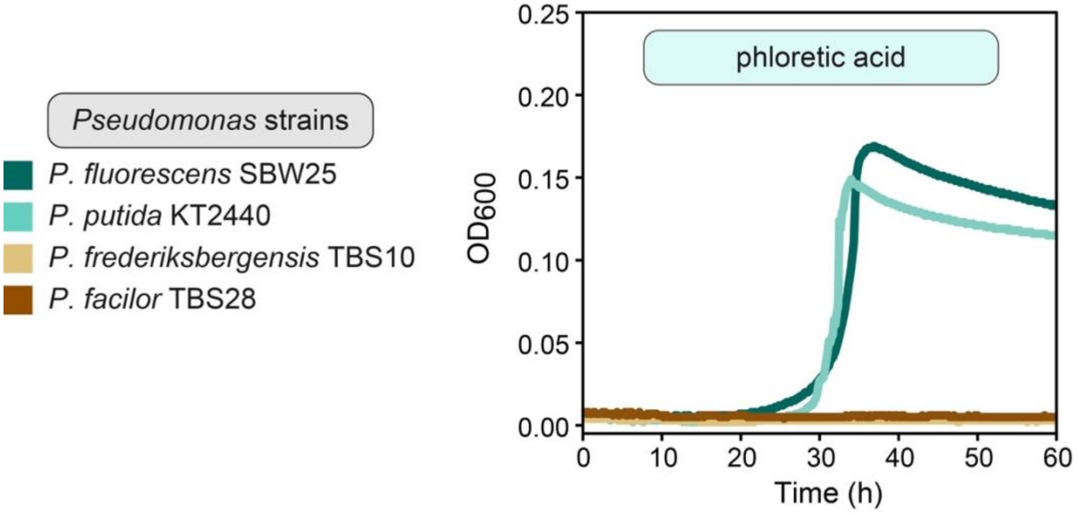


**Supplementary Fig. S2. A subset of environmental Pseudomonas isolates can utilize phenylpropanoid compound phloretic acid as a sole carbon source.** Microtiter plate cultivation assays comparing growth of four environmental Pseudomonads in MME medium containing 2.5 mM phloretic acid as the sole carbon source. Panel contains a single representative curve from one of three biological replicates.

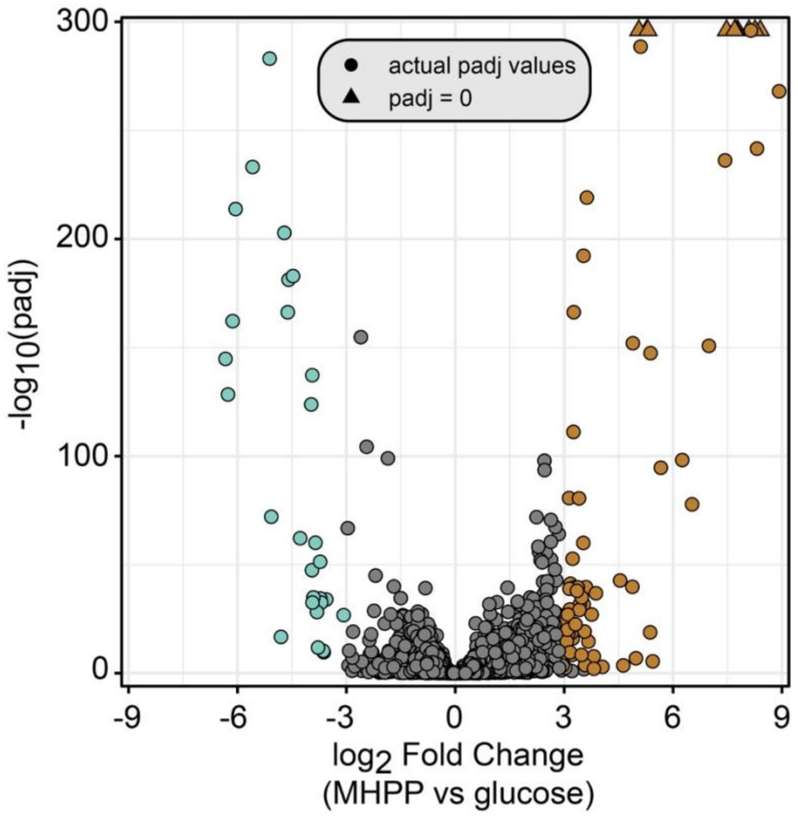


**Supplementary Fig. S3. Chart of differential gene expression when *P. fluorescens* SBW25 was grown with either glucose or MHPP as a carbon source.** Positive values on the X-axis indicate genes whose expression was higher when MHPP was the carbon source than when glucose was the carbon source. The y-axis represents a log transformation of the p-value that has been adjust for multiple tests. Brown and teal colored dots indicate genes whose differential expression was statistically significant and greater than 8-fold in crease in expression during growth on MHPP (brown) or glucose (teal). Samples with an adjusted p-value of 0 are shown at the top of the Y-axis as triangles, as they cannot be plotted otherwise.

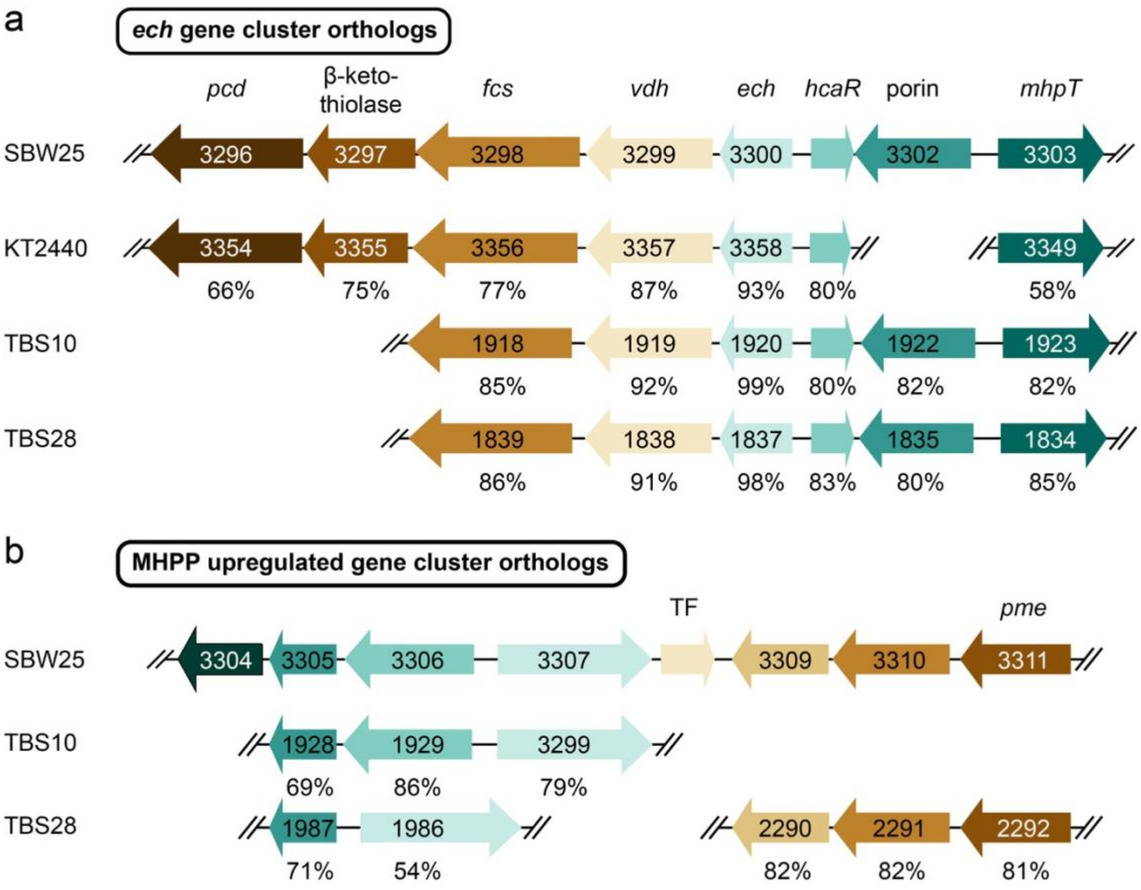


**Supplementary Fig. S4. Alignments of differentially expressed gene clusters containing genes critical for growth on MHPP and other phenylpropanoids in SBW25 with orthologs from other environmental Pseudomonads.** Locus tag numbers for genes in each organism are indicated in the arrows, with the exception of locus tags for *hcaR* and PFLU3308. Gene arrangement and spacing is maintained in each organism. Coloring in the arrows indicates orthologous genes. Percentage values under each arrow indicate the % identity of the SBW25 gene with the closest matching genes in KT2440, TBS10, or TBS28 gene. Genes with less than 50% identity were not considered. In organisms other than SBW25 genes from the *ech* gene cluster (a) are not co-localized with their equivalents of the MHPP upregulated gene cluster (b).

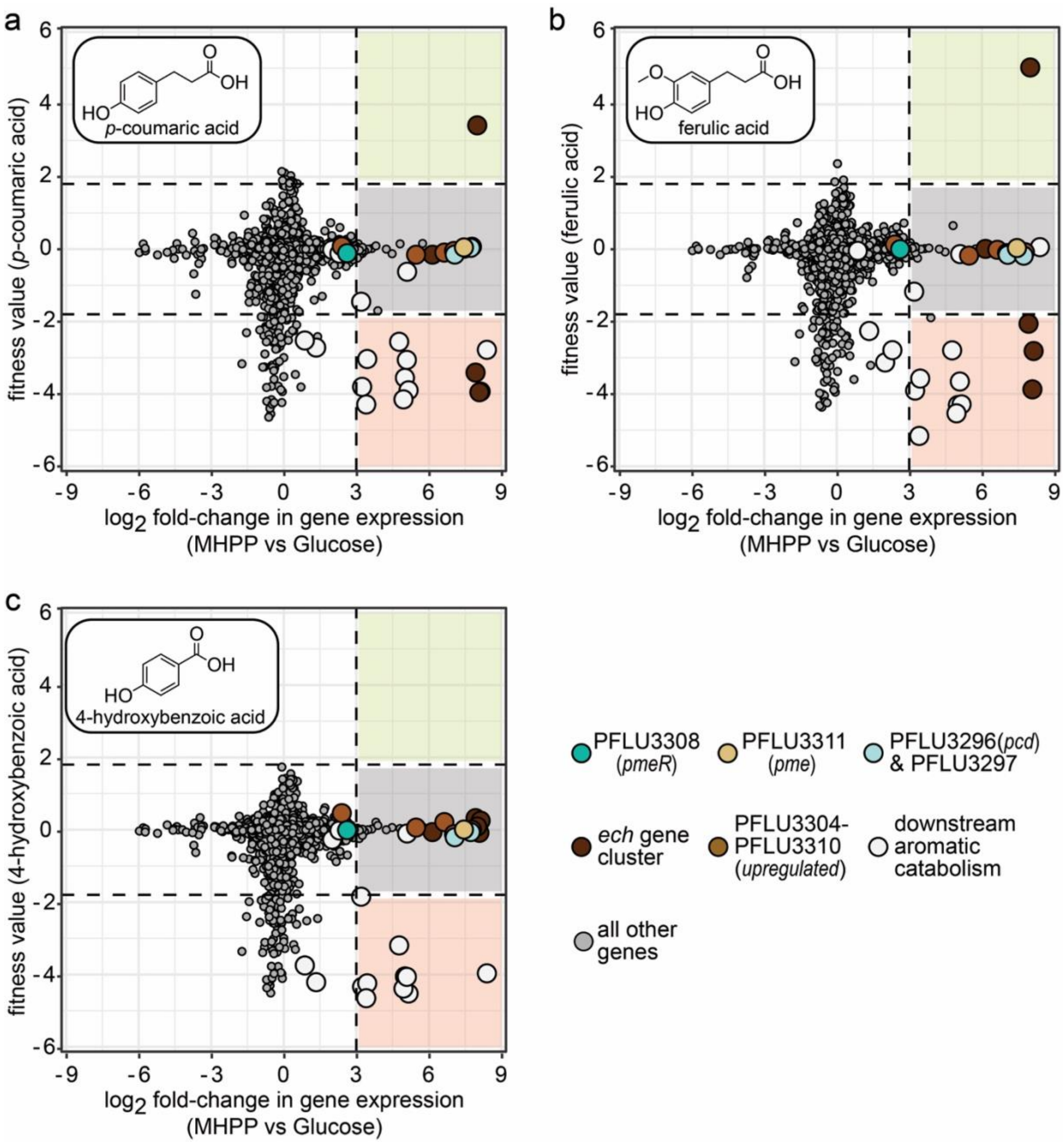


**Supplementary Fig. S5. Comparing fitness values with differential gene expression reduces search space for novel pathway genes.** (a-c) Plots comparing differential expression (x-axis) versus mean RB-TnSeq fitness values on the y-axis. RB-TnSeq values displayed are from cultures grown with either (a) *p*-coumaric acid, (b) ferulic acid, or (c) 4-hydroxybenzoic acid (a downstream metabolite of *p*-coumaric acid catabolism). Positive and negative differential expression values indicate higher expression during growth using MHPP and glucose as carbon sources, respectively. Dots indicate genes encoding the putative PPME-sensitive *pmeR* transcription factor (dark teal), phenylpropanoid methyl esterase (light brown), putative phloretoyl-CoA dehydrogenase and putative β-ketothiolase (light teal), other genes in the *ech* gene cluster (dark brown), other genes in the MHPP-upregulated gene cluster (medium brown), downstream aromatic catabolic pathway gene clusters (white), and all other genes (dark gray). Values represent the mean of 4 RB-TnSeq or 4 differential expression biological replicates. Genes lacking fitness or differential expression values are not displayed.

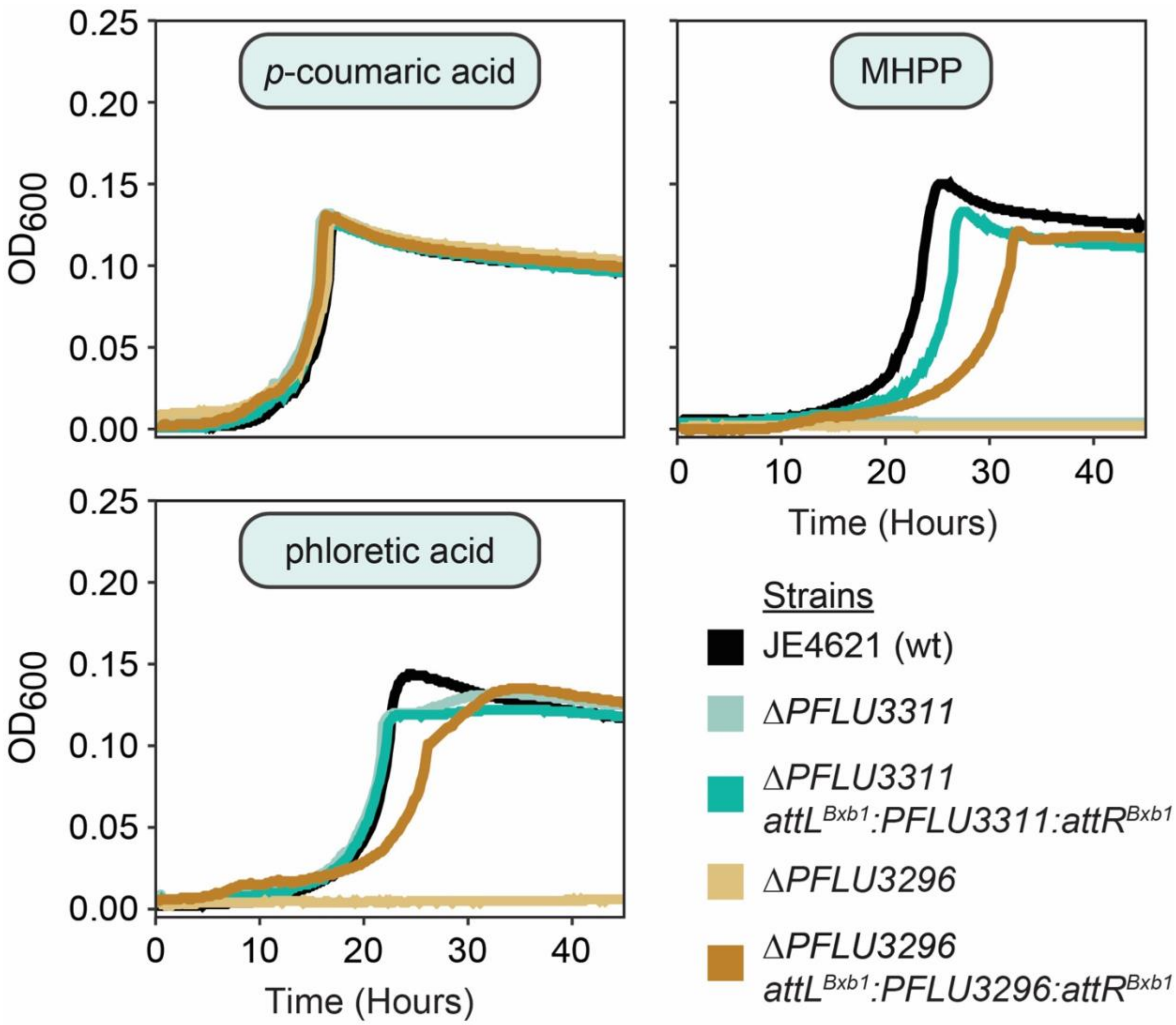


**Supplementary Fig. S6. Complementation of PFLU3296 and PFLU3311 gene deletions enables growth with MHPP and phloretic acid carbon sources.** Microtiter plate cultivation assays comparing growth of five *P. fluorescens* strains in MME medium containing 2.5 mM *p*-coumaric acid, 2.5 mM MHPP, or 2.5 mM phloretic acid as the sole carbon source. Each panel contains a single representative curve from one of three biological replicates.

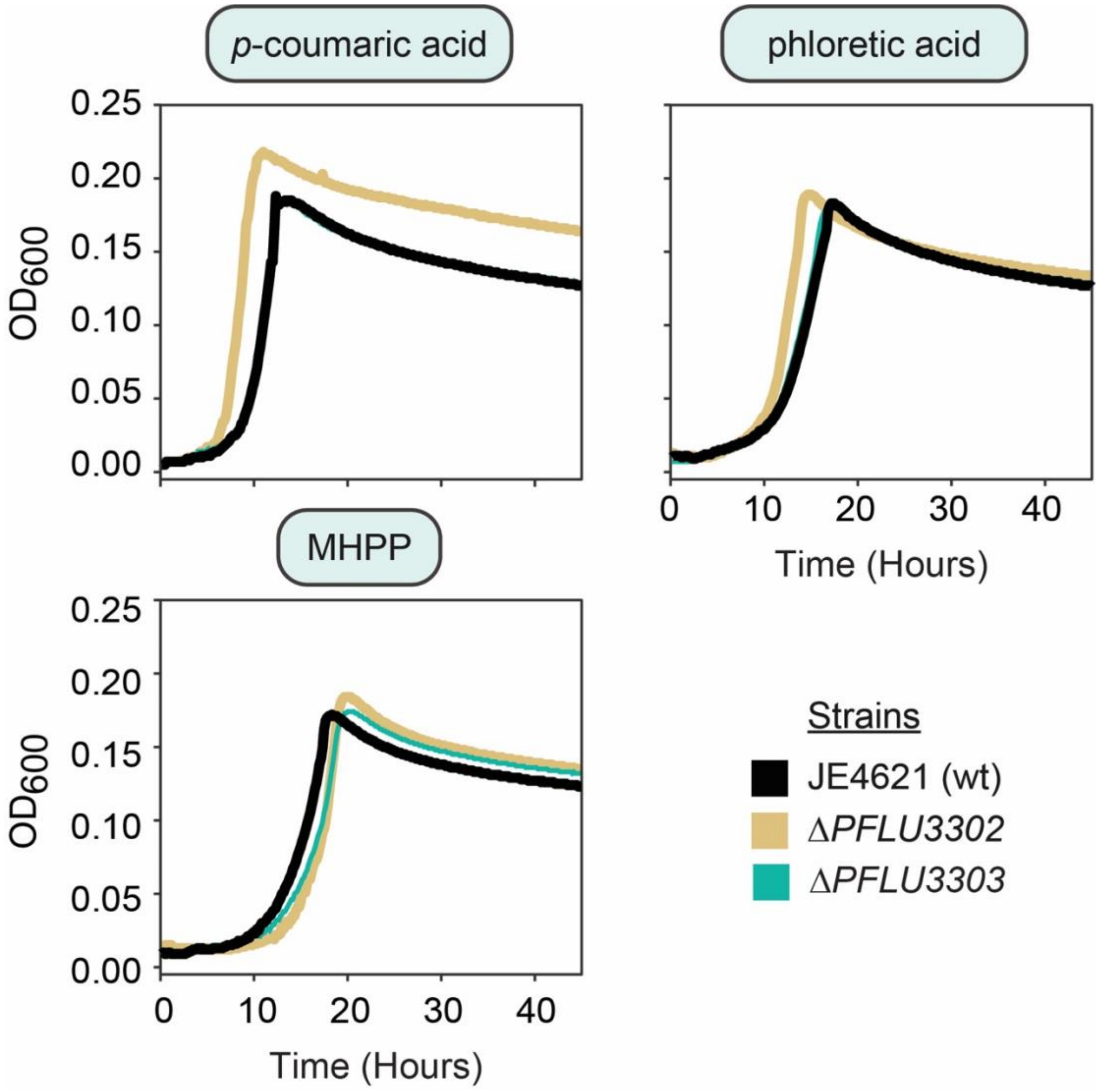


**Supplemental Fig. S7. Deletion of putative aromatic compound transporters has no apparent impact upon growth when using phenylpropanoids as carbon sources.** Microtiter plate cultivation assays comparing growth of three *P. fluorescens* strains in MME medium containing 2.5 mM *p*-coumaric acid, 2.5 mM MHPP, or 2.5 mM phloretic acid as the sole carbon source. Each panel contains a single representative curve from one of three biological replicates.

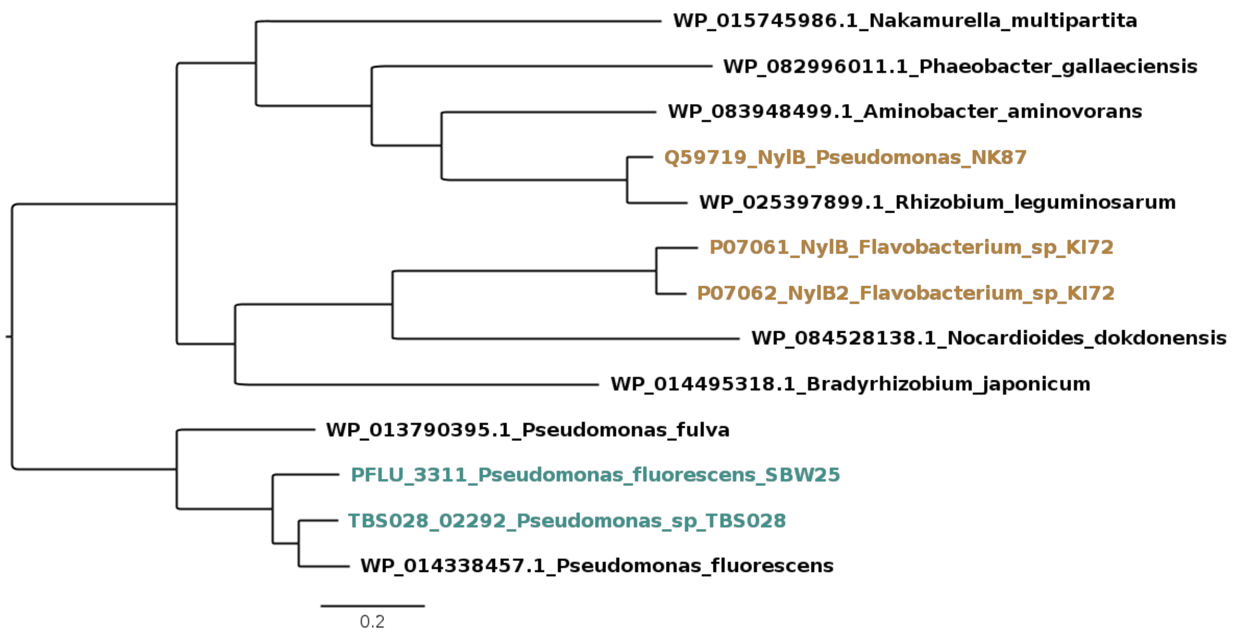


**Supplementary Fig. S8. Phylogenetic tree of serine hydrolases**. Serine hydrolase sequences from *P. fluorescens* SBW25 and *Pseudomonas facilor* TBS028 (teal) were aligned against proteins with known 6-aminohexanoate-dimer hydrolase activity (tan) and a selection of uncharacterized proteins with the closest sequence similarity to the methyl esterases (black). The proteins of interest only share ~35% AAID with the KI72 NylB and NylB2 sequences[1,2].

**Supplementary Table S1. Orthologs of SBW25 genes in other Pseudomonads**

| | | | KT2440 | | | | TBS10 | | | | TBS28 | | | |
|---|---|---|---|---|---|---|---|---|---|---|---|---|---|---|
| | SBW25 locus | product | locus | % identity | % positive | % gaps | locus | % identity | % positive | % gaps | locus | % identity | % positive | % gaps |
| ech operon | PFLU3296 | acyl-CoA dehydrogenase | PP3354 | 66 | 74 | 2 | --- | --- | --- | --- | --- | --- | --- | --- |
| | PFLU3297 | thiolase family protein | PP3355 | 75 | 81 | 0 | --- | --- | --- | --- | --- | --- | --- | --- |
| | PFLU3298 | feruloyl-CoA synthase | PP3356 | 77 | 85 | 0 | PFR10_01918 | 85 | 92 | 0 | PFA28_01839 | 86 | 92 | 0 |
| | PFLU3299 | aldehyde dehydrogenase | PP3357 | 87 | 92 | 0 | PFR10_01919 | 92 | 94 | 0 | PFA28_01838 | 91 | 94 | 0 |
| | PFLU3300 | p-hydroxycinnamoyl CoA hydratase/lyase | PP3358 | 93 | 96 | 0 | PFR10_01920 | 99 | 100 | 0 | PFA28_01837 | 98 | 99 | 0 |
| ech regulator | PFLU3301 | MarR family transcriptional regulator | PP3359 | 80 | 87 | 0 | PFR10_01921 | 80 | 88 | 0 | PFA28_01836 | 83 | 90 | 0 |
| conserved PP transporters | PFLU3302 | OprD family porin | --- | --- | --- | --- | PFR10_01922 | 82 | 90 | 0 | PFA28_01835 | 80 | 89 | 0 |
| | PFLU3303 | 3-(3-hydroxy-phenyl)propionate transporter MhpT | PP3349 | 58 | 72 | 0 | PFR10_01923 | 82 | 89 | 0 | PFA28_01834 | 85 | 90 | 0 |
| uncharacterized MHPP-upregulated gene cluster | PFLU3304 | hypothetical protein | --- | --- | --- | --- | --- | --- | --- | --- | --- | --- | --- | --- |
| | PFLU3305 | coniferyl alcohol dehydrogenase | --- | --- | --- | --- | PFR10_01928 | 69 | 78 | 0 | PFA28_01987 | 71 | 81 | 0 |
| | PFLU3306 | benzaldehyde dehydrogenase | --- | --- | --- | --- | PFR10_01929 | 86 | 92 | 0 | --- | --- | --- | --- |
| | PFLU3307 | sigma-54-dependent Fis family transcriptional regulator | --- | --- | --- | --- | PFR10_01930 | 79 | 85 | 0 | PFA28_01986 | 54 | 69 | 4 |
| | PFLU3308 | TetR/AcrR family transcriptional regulator | --- | --- | --- | --- | --- | --- | --- | --- | --- | --- | --- | --- |
| | PFLU3309 | polyamine ABC transporter substrate-binding protein | PP5341 | 55 | 73 | 0 | PFR10_04530 | 54 | 73 | 0 | PFA28_02290 | 82 | 90 | 0 |
| | PFLU3310 | hypothetical protein | --- | --- | --- | --- | --- | --- | --- | --- | PFA28_02291 | 82 | 88 | 0 |
| | PFLU3311 | serine hydrolase | --- | --- | --- | --- | --- | --- | --- | --- | PFA28_02292 | 81 | 87 | 0 |

Results from BLASTp pairwise alignment of SBW25 proteins with most highly similar proteins in the indicated organism. No value is listed if the most similar protein has <50% identity and <90% query coverage.

| Supplementary Table S2. Sequences of functionally verified serine hydrolases. | |
|---|---|
| **Protein** | **Protein Sequence** |
| Pme from Pseudomonas fluorescens SBW25 (PFLU3311) | MGQNIMPSLASLYVETNESSTQPRLAPLLMQGFPAGPKYRVTWHNWMRPPFNQWGFRNLA<br>RLRPSIDVRAGAGPAGPLNTVSQALDALYFDSECGLSVSVIEHLLASQTDAFLVMQGDTV<br>LFERYFNGQRPCDRHIMFSVTKSLVGTLGEELVTRGVLNPELPAGYYVPELVGSAFGDAT<br>VRQLFDMAVGIDYSEVYDDPNSESSQYGYACGFQPALAQYAQFESLYQYLPSLKKRGVHG<br>GFFHYVTATTEALAWVMERASGSACSELLEGIWQQLGCDRDGYFIADPWGRNVAGAGFSA<br>TLRDMARFGRLLANNGRQDGVELLSPETVARITAGADPAVYAQNAEFSHWTPGASYRSQW<br>YVFNDHSQALMAGGIHGQYLFIDKPSGVVIVKQSSLNEAVSPFDTDSVRMLRAIAAHLSH |
| Pme from Pseudomonas facile TBS28 | MSQNAVPSLASLYVEAIESSDPRSTSSFMQGFPPEPQRRVSWHNWMRAPFNQWGFRNLAR<br>LRPSIDVQAGAAPVSPLQQAPQPLDQLHFNSECGLSISVIEHLLASQTDAFLVMQGDTVL<br>YERYFNGQRPQDRHIMFSVTKSLIGTLGEQLVCEGLLDTALPAAHYVPELAGSAFADATV<br>RQLFDMAVGIDYSEVYEDPDSESSQYGYACGFQPAPVQYGQFESLYEYLPSLRKRGSHGG<br>FFHYVTATTEALAWVMERACGRACHELLQDIWSQLGCERDGYFMADPWGRNVAGAGFSAT<br>LRDMARFGRLLANEGRHAGRQLLSSEAIAGILAGADPAVYATSPDFSAWTPGASYRSQWY<br>VFNDHSQALMAGGIHGQYLFVDKPSGVVIVKQSSLSEAVSPFDGDSVRMLRAIAAHLTR |
| NylB from Paenarthobacter ureafaciens KI72 (previously Arthobacter sp. KI72) | MNARSTGQHPARYPGAAAGEPTLDSWQEAPHNRWAFARLGELLPTAAVSRRDPATPAEPV<br>VRLDALATRLPDLEQRLEETCTDAFLVLRGSEVLAEYYRAGFAPDDRHLLMSVSKSLCGT<br>VVGALIDEGRIDPAQPVTEYVPELAGSVYDGPSVLQVLDMQISIDYNEDYVDPASEVQTH<br>DRSAGWRTRRDGDPADTYEFLTTLRGDGGTGEFQYCSANTDVLAWIVERVTGLRYVEALS<br>TYLWAKLDADRDATITVDQTGFGFANGGVSCTARDLARVGRMMLDGGVAPGGRVVSQGWV<br>ESVLAGGSREAMTDEGFTSAFPEGSYTRQWWCTGNERGNVSGIGIHGQNLWLDPRTDSVI<br>VKLSSWPDPDTRHWHGLQSGILLDVSRALDAV |
| NylB from Pseudomonas sp. NK87 | MNTVPPFRDPTVPGNSHIPRQDWDRAPWNRWTFQHVRELLPTTKVWRGTGPASPLPVDLR<br>DIDAVSFAAEGQSHTIAGFLETSYADGFLVLHGGKIVAERYLNGMAPHTQHLSQSVAKSV<br>VGTVAGILIDRGVVNPAALLTHYLPELEATAYRGATVQHVLDMTSGVVFDETYTALDSHM<br>AQLDVASGWKDSPNPDWPTHVWDLILSLKDLECPHGASFRYRSIETDVLAFVLQRAAAAP<br>LAELVSRELWAPMGAEEDAYFTVDTAGYALGDGGFNATLRDYARFALLHLRGGEIDGRRI<br>VSPGWIAATRFGADPALFGDIYREALPAGAYHNQFWIEDTARGAYMARGVFGQLIYIDPE<br>ADFAAVILSSWPEFVSTTRLRTALAAVRAVREALSA |

**Supplementary Table S3. Strains and Plasmids used in this study.**

| Name | Relevant Genotype | Source |
|---|---|---|
| *Strains* | | |
| NEB 5-alpha F'Iq | *Escherichia coli* F´ *proA*$^{+}$*B*$^{+}$ *lacI*$^{q}$ *Δ(lacZ)M15 zzf::Tn10* (Tet$^{R}$) / *fhuA2Δ(argF-lacZ)U169 phoA glnV44 Φ80Δ(lacZ)M15 gyrA96 recA1 relA1 endA1 thi-1 hsdR17* | New England Biolabs |
| SBW25 | *Pseudomonas fluorescens SBW25* | [3] |
| KT2440 | *Pseudomonas putida KT2440* | [4] |
| TBS10 | *Pseudomonas frederiksbergensis* TBS10 | [5] |
| TBS28 | *Pseudomonas facilor* TBS28 | this work |
| Pf-5 | *Pseudomonas protegens* Pf-5 | [6] |
| DSM4188 | *Pseudomonas stutzeri* DSM4188 | [7] |
| TBS49 | *Pseudomonas* sp. TBS49 | this work |
| JE4621 | *P. fluorescens* SBW25 3' *ampC*:poly-*attB* | [5] |
| JE90 | *P. putida* KT2440 ΔhsdR::Bxb1int-*attB* | [8] |
| RS175 | *P. facilor* 3' *ampC*:poly-*attB* | this work |
| JE5041 | *P. frederiksbergensis* TBS10 *att*$^{Tn5}$:10x poly-*attB*:*att*$^{Tn5}$ | this work |
| RS137 | *P. fluorescens* JE4621 ΔPFLU3296 (pcd) | this work |
| AW65 | *P. fluorescens* JE4621 ΔPFLU3297 | this work |
| AW54 | *P. fluorescens* JE4621 ΔPFLU3298 (*fcs*) | this work |
| AW51 | *P. fluorescens* JE4621 ΔPFLU3300 (*ech*) | this work |
| RS183 | *P. fluorescens* JE4621 ΔPFLU3302 | this work |
| RS184 | *P. fluorescens* JE4621 ΔPFLU3303 | this work |
| AF001 | *P. fluorescens* JE4621 ΔPFLU3311 (*pme*) | this work |
| RS137-AW30 | *P. fluorescens* JE4621 ΔPFLU3296 *attL*$^{Bxb1}$:pAW30:*attR*$^{Bxb1}$ | this work |
| AF001-JE1918 | *P. fluorescens* JE4621 ΔPFLU3311 *attL*$^{Bxb1}$:pJE1918:*attR*$^{Bxb1}$ | this work |
| AF001-JE1944 | *P. fluorescens* JE4621 ΔPFLU3311 *attL*$^{Bxb1}$:pJE1944:*attR*$^{Bxb1}$ | this work |
| JE5041-JE1920 | *P. fredericksbergensis* JE5041 *attL*$^{Bxb1}$:pJE1920:*attR*$^{Bxb1}$ | this work |
| JE90-JE1918 | *P. putida* JE90 *attL*$^{Bxb1}$:pJE1918:*attR*$^{Bxb1}$ | this work |
| RS175-AW30 | *P. facilor* RS175 *attL*$^{Bxb1}$:pAW30:*attR*$^{Bxb1}$ | this work |
| JE5041-JE1045 | *P. fredericksbergensis* JE5041 *attL*$^{Bxb1}$:pJE1045:*attR*$^{Bxb1}$ | this work |
| JE90-JE1045 | *P. putida* JE90 *attL*$^{Bxb1}$:pJE1045:*attR*$^{Bxb1}$ | this work |
| RS175-JE1045 | *P. facilor* RS175 *attL*$^{Bxb1}$:pJE1045:*attR*$^{Bxb1}$ | this work |
| *Plasmids* | | |
| pK18sB | pUC origin, *nptII*, *sacB* | [9] |
| pGW31 | pUC origin, AprR, P$_{tac}$:*Bxb1 integrase* ΔmSF$^{ts1}$ | [5] |
| pJE1045 | pJE990 P$_{tac}$:*mNeonGreen* | [10] |
| pGW60 | pJE990 P$_{tac\text{-}mod}$:*mNeonGreen,* 10x poly-*attP* cassette | [5] |
| pJE1918 | pJE990 P$_{tac}$:PFLU3311 | this work |
| pJE1920 | pJE990 P$_{tac}$:PFLU3311:PFLU3296 | this work |

| | | |
|---|---|---|
| pAW30 | pJE990 $P_{tac}$:PFLU3296 | this work |
| pRS329 | pK18sB 3'*ampC*:poly-*attB* in TBS28 | this work |
| pRS309 | pK18sB ΔPFLU3296 | this work |
| pAW5 | pK18sB ΔPFLU3297 | this work |
| pAW11 | pK18sB ΔPFLU3298 | this work |
| pAW6 | pK18sB ΔPFLU3300 | this work |
| pRS337 | pK18sB ΔPFLU3302 | this work |
| pRS338 | pK18sB ΔPFLU3303 | this work |
| pAF002 | pK18sB ΔPFLU3311 | this work |
| pEVF-SBP1-P2 | PFLU3311 expression vector | this work |
